# Functional Landscape of Zebrafish Gonadotropins and Receptors: A Comprehensive Genetic Analysis

**DOI:** 10.1101/2025.11.26.690707

**Authors:** Chu Zeng, Zhiwei Zhang, Lingling Zhou, Xianqing Zhou, Nana Ai, Wei Ge

## Abstract

In vertebrates, reproduction is controlled by two master hormones, or gonadotropins, from the pituitary: follicle-stimulating hormone (FSH) and luteinizing hormone (LH), which act through their cognate receptors, FSHR and LHCGR, in gonads. Like other vertebrates, zebrafish also has two gonadotropins and respective receptors. However, while zebrafish FSH activates Fshr specifically, its LH can activate both Fshr and Lhcgr, suggesting three signaling pathways – the canonical FSH-Fshr and LH-Lhcgr pathways, and a non-canonical LH-Fshr pathway. To dissect functional roles of these pathways, we generated a series of zebrafish mutants lacking one to all four genes encoding the two ligands and their receptors: *fshb*, *lhb*, *fshr*, and *lhcgr.* Single mutants confirmed the essential roles of FSH and LH in zebrafish reproduction. Double mutants demonstrated the functionality of all three pathways, especially the non-canonical LH-Fshr pathway, and highlighted the importance of Fshr in mediating both FSH and LH actions. Transcriptome analysis of double mutant follicles (infertile *lhb-/-;lhcgr-/-* with FSH-Fshr and fertile *fshb-/-;lhcgr-/-* with LH-Fshr) revealed potential molecular mechanisms underlying LH stimulation of oocyte maturation and ovulation via Fshr. Triple mutants revealed spontaneous, ligand-independent activities of Fshr and Lhcgr, in supporting spermatogenesis. Males with only *fshb* or *lhb* were infertile, with testes dominated by spermatogonia and spermatocytes with few spermatozoa. The quadruple mutant (*fshb-/-;lhb-/-;fshr-/-;lhcgr-/-*) displayed an all-male phenotype with underdeveloped, infertile gonads primarily containing spermatogonia, suggesting a critical role for gonadotropins in sex differentiation and gonadal development. This comprehensive genetic study provides insights into functional importance of gonadotropins in zebrafish reproduction and their signaling mechanisms.

**Author summary:** Follicle-stimulating hormone (FSH) and luteinizing hormone (LH) are two key hormones regulating vertebrate reproduction via their specific receptors, FSHR and LHCGR. Zebrafish, like other vertebrates, also has two hormones and two receptors. Interestingly, zebrafish LH signals not only through its cognate receptor Lhcgr but also through FSH receptor Fshr, suggesting existence of three signaling pathways (FSH-Fshr, LH-Fshr and LH-Lhcgr). To investigate the functionality of these pathways, we created various zebrafish mutant lines with all mutational combinations for these hormones and receptors. Our findings demonstrated that Fshr, when activated by either FSH or LH, played a critical role in ovarian growth and folliculogenesis. LH was necessary for oocyte maturation and ovulation, and the LH-Fshr pathway alone was adequate to support the entire process of female reproduction. All three pathways were sufficient to sustain male reproductive function. We further observed that the receptors exhibited low-level spontaneous activities in the absence of hormones, that could support spermatogenesis. Fish lacking all four genes developed only as males with extremely underdeveloped, non-functional testes. This highlights critical roles for gonadotropins not only in fertility, but also in sex differentiation. Our work provides comparative insights into the fundamental mechanisms by which gonadotropins control reproductive processes in vertebrates.

## Introduction

Vertebrate reproduction is primarily controlled by two gonadotropins from the pituitary, namely follicle-stimulating hormone (FSH) and luteinizing hormone (LH). As members of a conserved glycoprotein hormone family, gonadotropins play essential roles in gametogenesis and steroidogenesis in both males and females [1–3]. Like other glycoprotein hormones, FSH and LH are both dimeric proteins composed of a common α subunit and a distinct β subunit [2, 4]. These hormones initiate their intracellular signaling cascades by binding to respective G protein-coupled receptors (FSHR and LHCGR) in gonads (ovary and testis), which signal primarily through cyclic adenosine monophosphate (cAMP) from fish to mammals [5–8]. In females, FSH binds to FSHR on granulosa cells to stimulate follicle growth and estrogen synthesis [9–11], while LH promotes androgen production in theca cells and initiates oocyte maturation and ovulation [12, 13]. In males, FSH acts on Sertoli cells to support sperm production and maturation, whereas LH stimulates testosterone production in Leydig cells [14–17].

Gene editing technology has been instrumental in elucidating the roles and functional importance of gonadotropin signaling in rodents. FSH or FSHR deficiency in female mice results in blocked folliculogenesis and infertility. Male mutants exhibit reduced testes size and fecundity [9, 18–21]. Similarly, LH or LHCGR deficiency in females leads to arrested folliculogenesis, while mutant males are infertile due to arrested spermatogenesis [14, 22, 23]. These models clearly demonstrate the critical importance of gonadotropin signaling in mammalian reproduction.

Unlike the well-defined receptor specificities observed in mammals, fish gonadotropins often exhibit promiscuous receptor recognition, making it challenging to dissect the specific roles of these hormones. For instance, in Japanese medaka (*Oryzias laptipes*), FSH and LH were reported to activate their respective receptors, Fshr and Lhcgr, but each hormone could also partially activate the other receptor [24]. However, this study should be interpreted with caution, as the hormones administered were recombinant single-chain polypeptides rather than natural dimeric proteins. In amago salmon (*Oncorhynchus rhodurus*), Lhcgr is primarily activated by LH but also shows slight responsiveness to FSH, whereas Fshr responds exclusively to FSH [25, 26]. In coho salmon (*Oncorhynchus kisutch*), Fshr does not differentiate between FSH and LH binding, whereas Lhcgr binds solely to LH [27]. In Atlantic salmon (*Salmo salar* L.), Fshr and Lhcgr also showed similar responses to purified coho salmon FSH and LH with LH also stimulating Fshr at 6-fold higher concentrations [28]. Similarly, in African catfish (*Clarias gariepinus*), Senegalese sole (*Solea senegalensis*) and common carp (*Cyprinus carpio*), Fshr is responsive to both recombinant FSH and LH, while Lhcgr is specifically activated by LH only [29–32].

In zebrafish (*Danio rerio*), a model fish species for studying development and reproduction, recombinant FSH and LH can activate their respective receptors, Fshr and Lhcgr; however, LH can also activate Fshr at high concentrations [33, 34]. This pattern is similar to observations in coho salmon, African catfish, Senegalese sole and common carp. These findings suggest the existence of three potential signaling pathways: two canonical pathways (FSH-Fshr and LH-Lhcgr) and one non-canonical pathway (LH-Fshr) [34]. However, due to technical constraints, there has been lack of studies specifically addressing the functionality of these pathways.

Recent genetic studies in zebrafish have provided valuable insights into the individual roles of two gonadotropins (FSH and LH) and their receptors (Fshr and Lhcgr) in regulating gonadal development and function in both sexes, further supporting the notion of promiscuous hormone-receptor interactions. Disruption of *fshb* gene in zebrafish significantly delayed the onset of folliculogenesis in the ovary and spermatogenesis in the testis. However, FSH does not appear to be essential for zebrafish reproduction, as its functions can be compensated at later stage by LH, which is also capable of binding and activating Fshr at higher concentrations [35]. This phenotype differs from the complete block of folliculogenesis prior to antral follicle formation and infertility in FSH-deficient mice [9]. In contrast, knockout of the *fshr* gene in zebrafish resulted in a complete arrest of follicle development at the early primary growth (PG) stage, indicating its indispensable role in early folliculogenesis [36]. Given that Fshr can be activated by both FSH and LH, it would be intriguing to distinguish the functional differences between the canonical FSH-Fshr pathway and the non-canonical LH-Fshr pathway. Interestingly, *lhcgr* knockout zebrafish showed no obvious phenotypes, with both male and female mutants exhibiting normal sex differentiation, puberty onset, gonadal development, and fertility [36]. These findings suggest that LH may exert its effects through alternative pathways, most likely involving Fshr [35, 37]. This raises an interesting question about the functionality of the canonical LH-Lhcgr pathway in zebrafish reproduction.

An intriguing aspect of gonadotropin signaling is the phenomenon of ligand-independent, constitutive receptor activity [38, 39]. In mammals, the constitutive activity of FSHR has been proposed to support Leydig cell function in FSH-deficient mice [40]. Similarly, in teleosts, Lhcgr has been suggested to exhibit ligand-independent activity in vitro [33, 41, 42]. Our previous studies on double mutants lacking both FSH and LH ligands showed normal spermatogenesis in male zebrafish despite much smaller testes, implying a role for constitutive receptor activity of gonadotropin receptors in the absence of ligands [35]. This hypothesis was further supported by the impaired spermatogenesis and male infertility observed in the double receptor mutant *fshr*-/-*;lhcgr-/-* [36]. However, the precise contributions of constitutive activity from individual receptors, Fshr and Lhcgr, remain to be clarified.

To further investigate the roles of gonadotropins and their receptors in zebrafish gonadal development and function, particularly the functionality of the three pathways (FSH-Fshr, LH-Lhcgr, and LH-Fshr) and the spontaneous activities of receptors Fshr and Lhcgr in both sexes, we conducted a comprehensive genetic analysis in this study. Our experiments involved generating all possible combinations of mutants for the four gonadotrophic genes involved (ligands *fshb* and *lhb,* and receptors *fshr* and *lhcgr*), including single, double, triple, and quadruple mutants. Phenotypic analyses of these mutants provided critical insights into the functional redundancy, compensatory mechanisms, and ligand-independent receptor activities of gonadotropin signaling in zebrafish. In addition, we also performed a transcriptome analysis on full-grown (FG) follicles from the fertile double mutants *fshb-/-;lhcgr-/-* with LH-Fshr and infertile *lhb-/-;lhcgr-/-* with FSH-Fshr to understand the molecular mechanisms underlying the roles of LH-Fshr and FSH-Fshr pathways in controlling final oocyte maturation and ovulation.

## Materials and Methods

### Zebrafish maintenance and husbandry

The mutant zebrafish lines *fshb* (ZFIN line No.: umo1; -10-bp deletion), *lhb* (umo2; - 5-bp deletion), *fshr* (umo3; -11-bp deletion), and *lhcgr* (umo4; -14-bp deletion) were generated in our previous studies [35, 36]. The adult fish were maintained under controlled conditions in the ZebTEC Multilinking Zebrafish system (Tecniplast, Buguggiate, Italy), with a temperature of 28 ± 1 °C, a pH of 7.3, and a photoperiod of 14-h light and 10-h darkness (light on at 8:30 am). From 5 to 10 days post-fertilization (dpf), the larval fish were fed paramecia, supplemented with egg yolk or Z Plus Premium Dry Larval Diet (Zeigler, Gardners, PA). From 10 to 15 dpf, their diet was switched to artemia. The fish were transferred to the main aquatic system at 15 dpf. They were fed with Otohime fish diet (Marubeni Nisshin Feed, Tokyo, Japan) twice daily using the Tritone automatic feeder system (Tecniplast), and additional artemia as a supplement. The animals were handled in accordance with the Animal Protection Act enacted by the Legislative Council of Macao Special Administrative Region under Article 71(1) of the Basic Law. All experimental protocols were approved by the Research Ethics Panel of the University of Macau (AEC-13-002).

### Genotyping

Genomic DNA from a small piece of the caudal fin from each individual was extracted by alkaline lysis [35, 43]. Briefly, 40 µL NaOH (50 mM) was added to each sample to extract the genomic DNA at 95°C for 10 min. The reaction was then neutralized by adding 8 µL of Tris-HCl (pH 8.0). The extracted genomic DNA was analyzed using High-Resolution Melt Analysis (HRMA) with the Precision Melt Analysis software (Bio-Rad Laboratories, Hercules, CA) to identify different genotypes based on melt curve differences [44]. The primers used for HRMA are listed in Table S1.

### Fish sampling and histological analysis

The fish were anaesthetized using MS222 (Sigma-Aldrich, Louis, MO) and photographed with a digital camera (EOS700D, Canon, Tokyo, Japan) to record gross morphology prior to processing. The body length (BL) and body weight (BW) of each fish were measured, and the gonads were photographed in situ after opening the abdominal cavity (Fig. 1A). During dissection, the addition of Bouin’s solution enhanced the visibility of the small testes. For histological analysis, the entire fish was fixed in Bouin’s solution for at least 24 h at room temperature. The fixed samples were dehydrated and infiltrated using the ASP6025S Automatic Tissue Processor (Leica, Wetzlar, Germany). The processed samples were embedded in paraffin followed by serial sectioning at a thickness of 5 μm. Hematoxylin and eosin staining (H&E) and slide mounting were performed using the ST5020 Stainer Integrated Workstation and the CV5030 Glass Coverslipper (Leica Biosystems). The sections were examined using the Eclipse Ni-U upright microscope (Nikon, Tokyo, Japan), imaged with the Digital Sight DS-i2 camera (Nikon), and assessed for sex ratio and mature spermatozoa (Fig. 1A).

**Fig 1.**
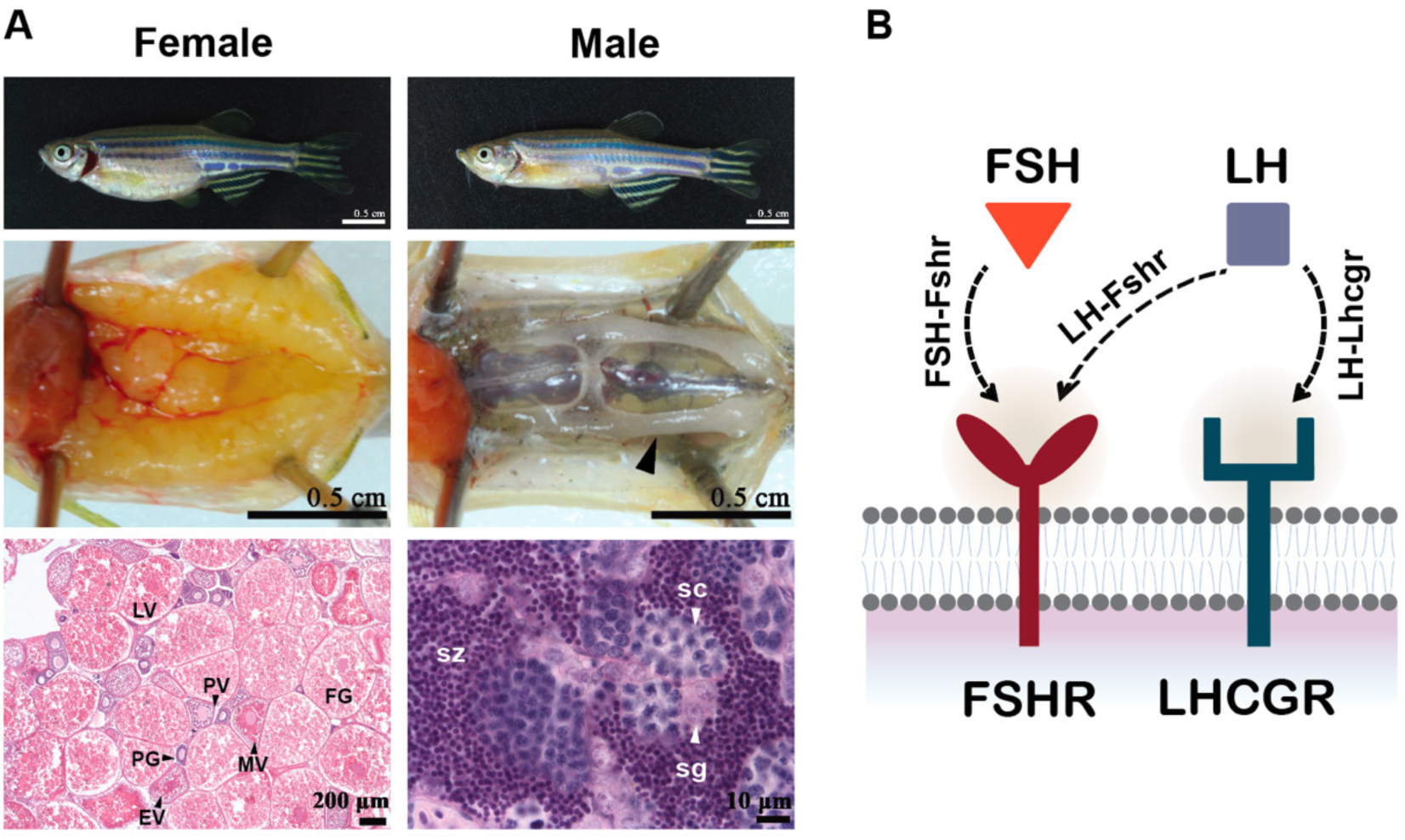
Overview of zebrafish gonadal structure and potential gonadotropin signaling pathways. (A) Representative images showing the overall body morphology, gonadal appearance, and histology of female and male zebrafish. (B) Diagram summarizing three possible gonadotropin signaling pathways.

### Determination and analysis of follicle composition

The follicle stages on sections were classified based on follicle size and morphological characteristics (cortical alveoli, yolk granules, etc.) as previously reported [45]. The follicles were classified into the following six stages [45]: PG (primary growth or stage I; <150 μm, lacking cortical alveoli), PV (previtellogenic or stage II; ∼250 μm, containing cortical alveoli but not yolk granules), EV (early vitellogenic or early stage III; ∼350 μm, containing both cortical alveoli and yolk granules), MV (mid-vitellogenic or mid-stage III, ∼450 μm), LV (late-vitellogenic or late stage III, ∼550 μm) and FG (full-grown, ∼650 μm). For follicle composition analysis, the entire fish was serially sectioned longitudinally at a thickness of 5 μm. The diameters of follicles were measured on three sections spaced at least 60 μm apart using NIS-Elements BR software (Nikon) [44]. Among these sections, the middle one was typically the largest in area. To ensure accurate measurement of follicle diameters, only follicles with visible nuclei (germinal vesicles) in the selected sections were measured.

### Measurement of testis size and quantification of mature spermatozoa

Testis size was measured by outlining and calculating the area of the three largest adjacent longitudinal sections from a single testis of each fish using ImageJ software. The average of three measurements per sample was used as the final testis size. Mature spermatozoa were identified based on their distinctive staining intensity and compact cellular structure. The areas occupied by mature spermatozoa within testicular sections were quantified using the Wood Tool in ImageJ, with the tolerance parameter set between 30 and 70. The proportion of area occupied by mature sperm relative to the total testis section area was used as an indicator of sperm production levels [46].

### Assessment of fertility and fecundity

Fish fertility and fecundity of different genotypes were assessed by natural mating with fertile wild-type (WT) partners. Three mutant fish were paired with three WT fish of the opposite sex in the spawning box (Tecniplast) at least three times at 3-day intervals, and the data presented represent the average of multiple tests performed on the three fish. For male mutants, the numbers of fertilized eggs from the WT female partner were counted, while for female mutants, total egg numbers were recorded including both fertilized and unfertilized ones. Individuals that failed to spawn or produce fertilized embryos after at least 5 trials were classified as infertile.

### RNA extraction and quantitative real-time PCR

Total RNA was isolated from the FG follicles using TRIzol (Invitrogen, Carlsbad, CA) according to the manufacture’s protocol, and quantified using the NanoDrop 2000 (Thermo Scientific, Waltham, MA). Reverse transcription reaction was performed using M-MLV reverse transcriptase (Invitrogen). Real-time qPCR was performed on the CFX96 Real-Time PCR Detection System using the SsoFast EvaGreen Supermix (Bio-Rad). Each sample was assayed in duplicate for accuracy, and the primers used are listed in Table S1. The expression of target genes in each sample was normalized to that of the housekeeping gene *ef1a*, and expressed as the fold change over the control group.

### Transcriptome analysis

To investigate the underlying mechanisms of LH-induced oocyte maturation and ovulation via Fshr, we collected FG follicles from 11 samples at 180 days post-fertilization (dpf) as follows: three control fish with all three pathways (FSH-Fshr, LH-Fshr, and LH-Lhcgr), four *lhb*-/-*;lhcgr*-/*-* fish with FSH-Fshr, and four *fshb*-/-*;lhcgr*-/-fish with LH-Fshr. Their ovaries were sampled between 3:30 and 4:30 am (4-5 hours before light on) as previous studies showed that the expression of *lhb* in the pituitary started to rise at 1:00 am (7 hours before light on) and remained high until 7:00 am [34, 47]. The sampled ovaries from each fish were placed in a microtube containing 1 mL L15 medium (Sigma-Aldrich). Fat tissues and ligaments surrounding the ovaries were removed using the BD Microlance 26G needle (BD, San Diego, CA). The ovarian fragments were then gently pipetted multiple times with a transfer pipette (JET Biofil, Guangzhou, China) to disperse the follicles. FG follicles (> 650 μm) were enriched by passing the suspended follicles through a 600 μm filter (SPL Lifesciences, Waunakee, WI) and manually selected for subsequent transcriptome (RNA-seq) analysis. The RNA extraction, sequencing, and preliminary analysis were carried out by Novogene Bioinformatics Technology (Tianjin, China). Briefly, the mRNA was purified from total RNA using poly-T oligo-attached magnetic beads and the sequencing libraries were prepared using the TruSeq RNA Library Prep Kit (Illumina, San Diego, CA). The libraries were sequenced on the Illumina Novaseq platform and 150 bp paired-end reads were generated. Differential expression analysis was performed using the DESeq2 R package (1.20.0). The resulting P-values were adjusted using the Benjamini and Hochberg’s approach to control the false discovery rate. Genes with an adjusted P-value < 0.05 were considered differentially expressed genes (DEGs) with statistical significance. DEGs were then subject to Gene Ontology (GO) enrichment and Kyoto Encyclopedia of Genes and Genomes (KEGG) pathway enrichment analyses. The RNA-seq data can be accessed at NCBI under the BioProject: PRJNA1276432.

### Data analysis

All data used for statistical analysis were obtained from multiple independent experiments and/or biological repeats (n ≥ 3) and expressed as mean ± SEM. The data were analyzed by Student’s t-test or one-way ANOVA followed by a Tukey multiple comparison test using Prism (GraphPad, La Jolla, CA). The significance level was denoted as * for *p* < 0.05, ** for *p* < 0.01, and *** for *p* < 0.001, or indicated by different letters.

## Results

### Single mutants confirm sex-specific roles of gonadotropins and their receptors

As in mammals, zebrafish FSH and LH preferentially act on their cognate receptors (Fshr and Lhcgr) respectively, forming two canonical pathways (FSH-Fshr and LH-Lhcgr). However, zebrafish LH can also partially activate Fshr at high concentrations, suggesting the existence of an additional non-canonical pathway (LH-Fshr) (Fig. 1B) as we proposed previously [34]. Recent studies in zebrafish on single mutants of gonadotropins and their receptors have established their functional roles in gametogenesis and reproduction [35–37, 48]. We also analyzed all single mutants (*fshb-/-*, *lhb-/-*, *fshr-/-*, and *lhcgr-/-*) in the present study to verify their phenotypes, which served as the references for other mutant combinations. Histological analysis at 120 dpf showed that all four single mutants exhibited normal testis development. In contrast, the mutant *fshb*-/- females displayed a significantly delayed ovarian growth, with follicle developing only to MV stage compared to age- and size-matched controls. In contrast, *lhb*-/- females exhibited normal ovarian growth with complete follicle development; however, they were unable to spawn eggs and, as a result, were infertile. As for receptors, *fshr*-/- females displayed extremely small ovaries, with all follicles arrested completely at early PG stage with large inter-follicular spaces. By comparison, *lhcgr-/-* females exhibited normal ovarian development and folliculogenesis with normal fertility (Fig. S1).

### Double mutants reveal functionality of ligand-receptor pathways and their cross-talks

To explore the ligand-receptor pathways and functional relationships among two gonadotropins and two receptors in zebrafish (Fig. 1B), we generated all six possible double mutants. Histological analysis indicated that the males of *fshb*-/-*;fshr*-/- with LH-Lhcgr pathway, *lhb*-/-*;lhcgr*-/- with FSH-Fshr pathway, and *fshb*-/-*;lhcgr*-/- with LH-Fshr pathway all showed normal testis structure with spermatogonia (sg), spermatocytes (sc), and spermatozoa (sz) at 90 dpf (Fig. 2A-D). However, the testis size of *fshb*-/-*;fshr*-/- was significantly lower than control and the double mutants lacking *lhcgr*, indicating an important role for Fshr in promoting testis growth (Fig. 3A). The level of sperm production as indicated by relative area of mature spermatozoa in the testis was also lower in *fshb*-/- mutants with either *fshr*-/- or *lhcgr*-/-, although these differences were not statistically significant compared to the control group, suggesting a role for FSH in promoting spermatogenesis (Fig. 3B). All three double mutants (*fshb*-/-*;fshr*-/-, *lhb*-/-*;lhcgr*-/- and *fshb*-/-*;lhcgr*-/-) could spawn normally with WT females, producing comparable number of fertilized eggs (Fig. 3C). Similar results were also observed at 360 dpf (Fig. 3D-E and Fig. S2A-D). These observations indicate that all three pathways (FSH-Fshr, LH-Fshr and LH-Lhcgr) were functional and each was sufficient to support testis growth, spermatogenesis and male reproduction in zebrafish.

**Fig 2.**
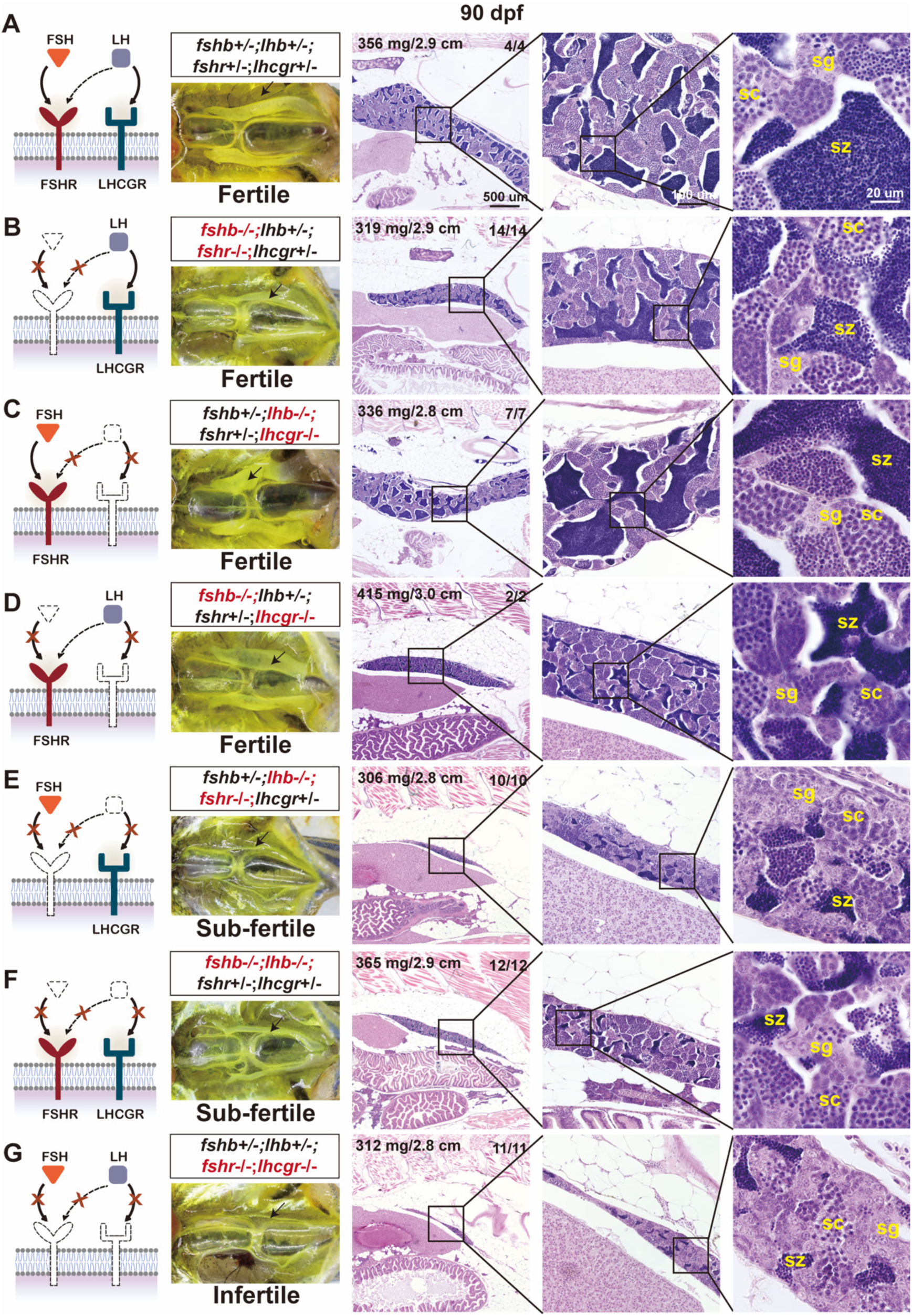
Histological examination of testicular development in double mutants at 90 dpf. (A) Representative control testes. (B-E) Functions of four potential pathways (LH-Lhcgr, FSH-Fshr, LH-Fshr and FSH-Lhcgr) in testis development. The pathways of LH-Lhcgr, FSH-Fshr and LH-Fshr could all support normal testis growth and spermatogenesis; however, the putative pathway FSH-Lhcgr did not appear to be functional in promoting normal spermatogenesis (small testis and limited spermatogenic activity). (F, G) Testis growth and spermatogenesis in double ligand mutants *fshb*-/-*;lhb*-/- (receptor-only) and double receptor mutants *fshr*-/*-;lhcgr*-/-(ligand-only). Both showed hypotrophic testis and poor spermatogenesis, especially the double receptor mutants. Black arrows point to the testes in abdominal cavity. The upper-left corner of the HE-stained image lists body weight (mg) and length (cm) of the fish, while the upper-right corner denotes the number of fish examined (lower) and the number showing similar phenotype (upper). sc, spermatocytes; sg, spermatogonia; sz, spermatozoa.

**Fig 3.**
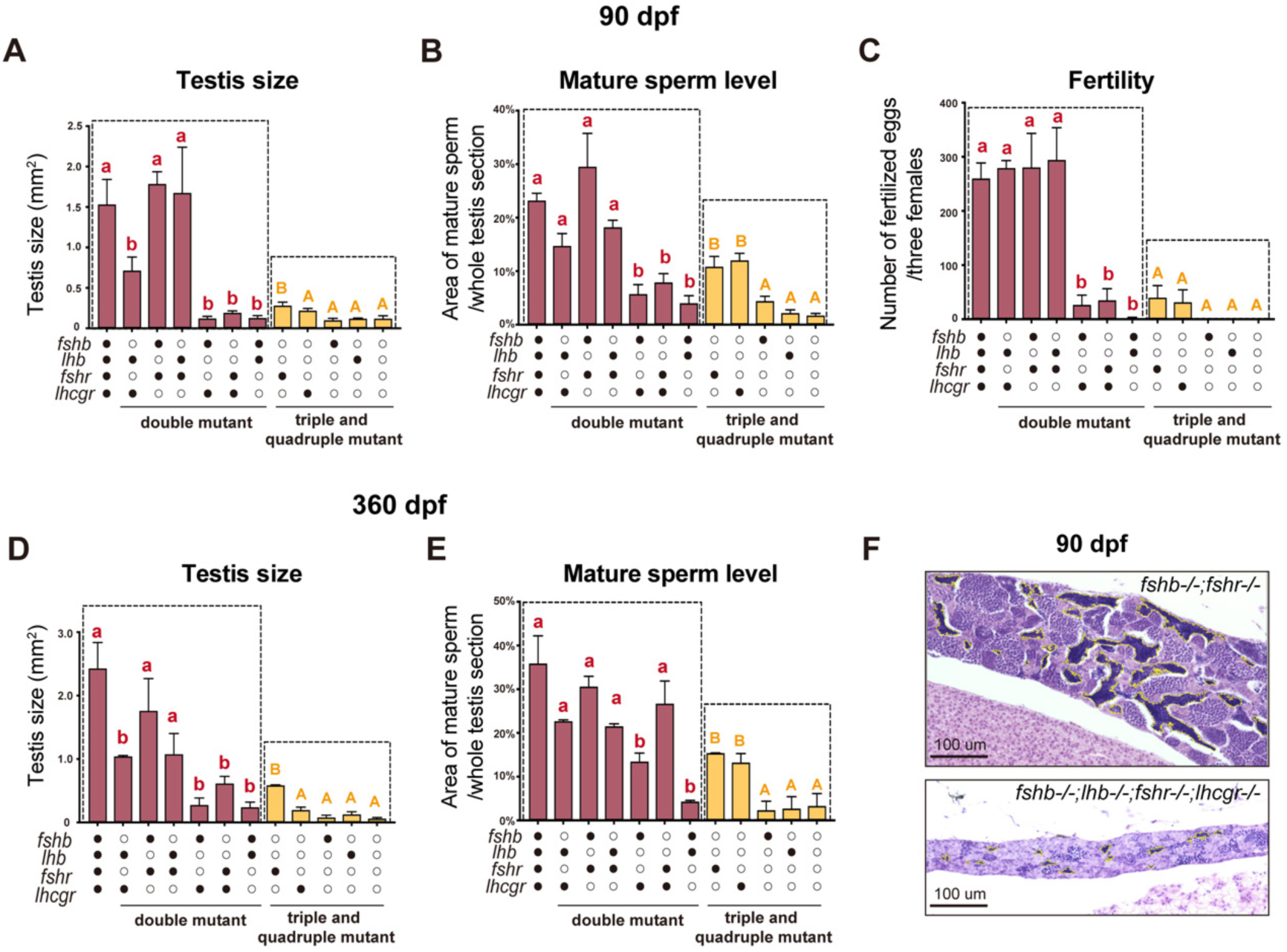
Quantitative assessment of testis size, mature sperm level and fertility in mutant males at 90 and 360 dpf. (A) Relative size of testes measured at 90 dpf. (B) Quantification of area proportion of mature sperm within the testes at 90 dpf. (C) Fertility test at 90 dpf. Individuals that failed to produce fertilized embryos after at least 5 independent trials were classified as infertile. (D) Relative size of testes measured at 360 dpf. (E) Quantification of area proportion of mature sperm within the testes at 360 dpf. All values were shown as means ± SEM and analyzed by one-way ANOVA (*p* < 0.05). Lowercase letters indicate significant differences from the control group, whereas uppercase letters mark significant differences from the quadruple mutant. ○: -/-; ●: +/-. (F) Schematic diagram illustrating the method used to quantify mature sperm production within the testis. Luminal regions containing mature spermatozoa were delineated with yellow lines and measured using ImageJ software.

Our previous study demonstrated a cross reaction between LH and Fshr, but not between FSH and Lhcgr, suggesting the existence of a non-canonical pathway LH-Fshr, but not FSH-Lhcgr in zebrafish [34]. This is further supported by the genetic data from the double mutant *lhb*-/-*;fshr*-/- with FSH and Lhcgr, which showed extremely underdeveloped testis, low level of sperm production, and poor fertility at both 90 and 360 dpf (Fig. 2E, Fig. S2E and Fig. 3), in contrast to the double mutant *fshb*-/-*;lhcgr*-/-with LH-Fshr, which showed full-grown testis, normal spermatogenesis, and normal fertility (Fig. 2D, Fig. S2D and Fig. 3). In addition, when comparing the triple mutant expressing *lhcgr* only (*fshb-/-;lhb-/-;fshr-/-;<u>lhcgr+/-</u>*) with the double mutant expressing both *fshb* and *lhcgr* (*<u>fshb+/-</u>;lhb-/-;fshr-/-;<u>lhcgr+/-</u>*), no improvements were observed in the three evaluated parameters (testis size, mature sperm production, and fertility) (Fig. 3 and Fig. 4). This result excludes the possibility that FSH can activate Lhcgr.

**Fig 4.**
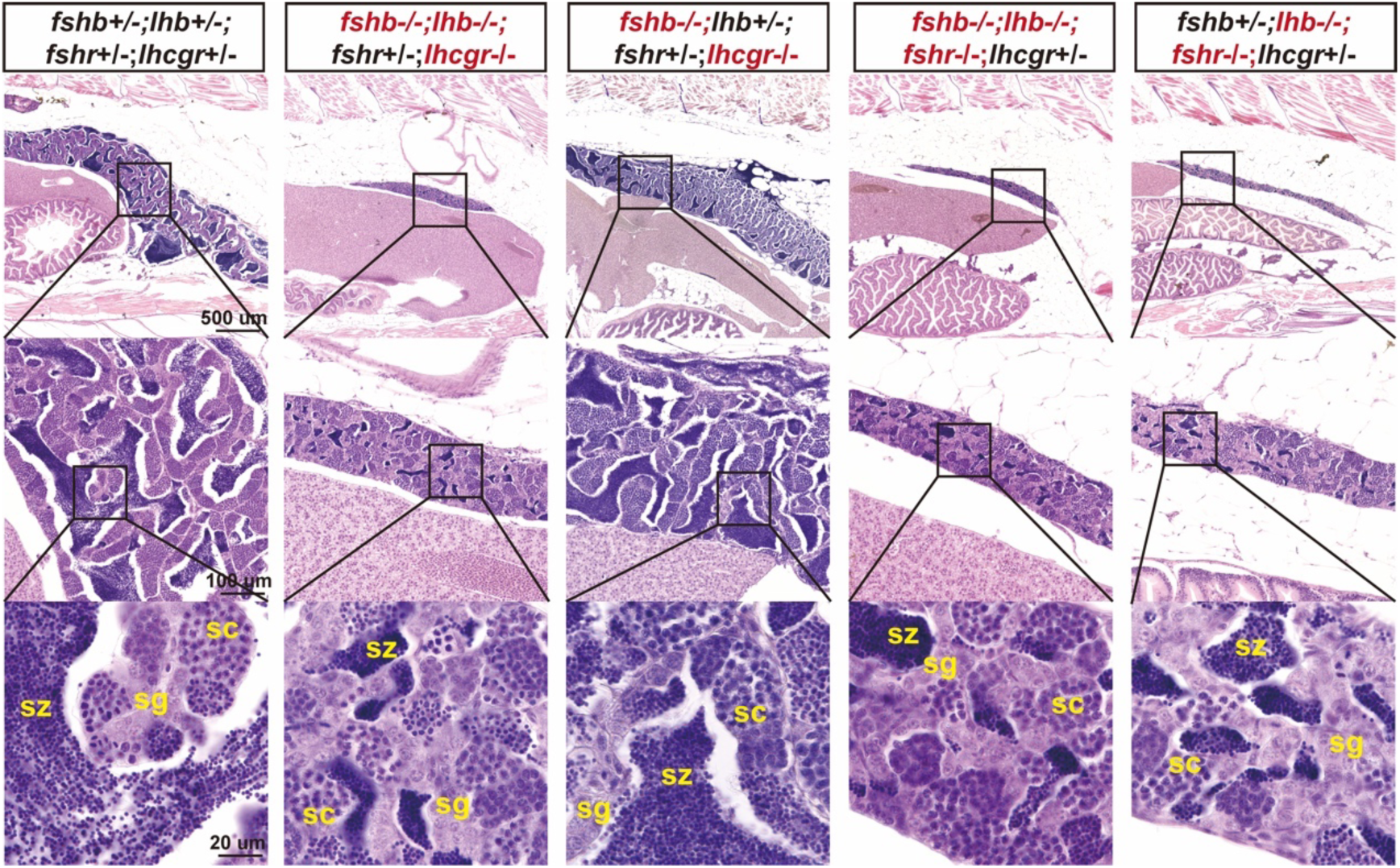
Histological evidence for LH cross-reaction with Fshr but not FSH with Lhcgr. Comparative histological analysis of the triple mutants expressing *fshr* only (*fshb-/-;lhb-/-;<u>fshr+/-</u>;lhcgr-/-*) and the double mutants expressing both *lhb* and *fshr* (*fshb-/-;<u>lhb+/-;fshr+/-</u>;lhcgr-/-*) showed potent stimulatory effect of LH on testis growth and spermatogenesis via Fshr. In contrast, analysis of the triple mutants expressing *lhcgr* only (*fshb-/-;lhb-/-;fshr-/-;<u>lhcgr+/-</u>*) and the double mutants expressing both *fshb* and *lhcgr* (*<u>fshb+/-</u>;lhb-/-;fshr-/-;<u>lhcgr+/-</u>*) revealed no effect of FSH on testis development via Lhcgr. sg, spermatogonia; sc, spermatocytes; sz, spermatozoa.

We also examined the phenotypes of double mutants lacking both ligands (*fshb*-/-*;lhb*-/-) or receptors (*fshr*-/-*;lhcgr*-/-). At 90 dpf, both double mutants showed similar phenotypes, including underdevelopment or hypotrophy of testis, low production of mature sperm, and poor fertility with double receptor mutant being completely infertile (Fig. 2F-G and Fig. 3). Interestingly, the testis size and sperm production bounced significantly at 360 dpf in the double ligand mutant (*fshb*-/-*;lhb*-/-); in contrast, the testes of double receptor mutant (*fshr-*/-*;lhcgr*-/-) remained extremely underdeveloped with germ cells being primarily spermatogonia (Fig. S2F-G and Fig. 3), indicating continual ligand-independent receptor activities.

We then analyzed ovarian development and folliculogenesis at 90 dpf in females of different double mutants, which provided critical insights into roles of the three pathways (FSH-Fshr, LH-Fshr, and LH-Lhcgr) in follicle development. The mutant females (*lhb*-/-*;lhcgr*-/-) with FSH-Fshr pathway showed normal ovarian growth and follicle composition (from PG to FG) compared to the control (Fig. 5A-B). In contrast, the *fshb*-/-*;lhcgr*-/- females with LH-Fshr pathway fish displayed a significant delay of follicle development as the mutant ovaries contained PG and PV follicles only without vitellogenic follicles, indicating that LH could act through Fshr, but the activation was not as strong as FSH (Fig. 5A-B). However, despite a delayed follicle development in young LH-Fshr fish at 90 dpf, its folliculogenesis eventually caught up and became fully comparable to that of control fish at 180 dpf (Fig. 5C). The most striking observation was the complete arrest of follicle development at early PG stage in *fshb*-/-*;fshr*-/- fish with LH-Lhcgr pathway, indicating no role for LH in driving folliculogenesis through its cognate receptor Lhcgr (Fig. 5A-B).

**Fig 5.**
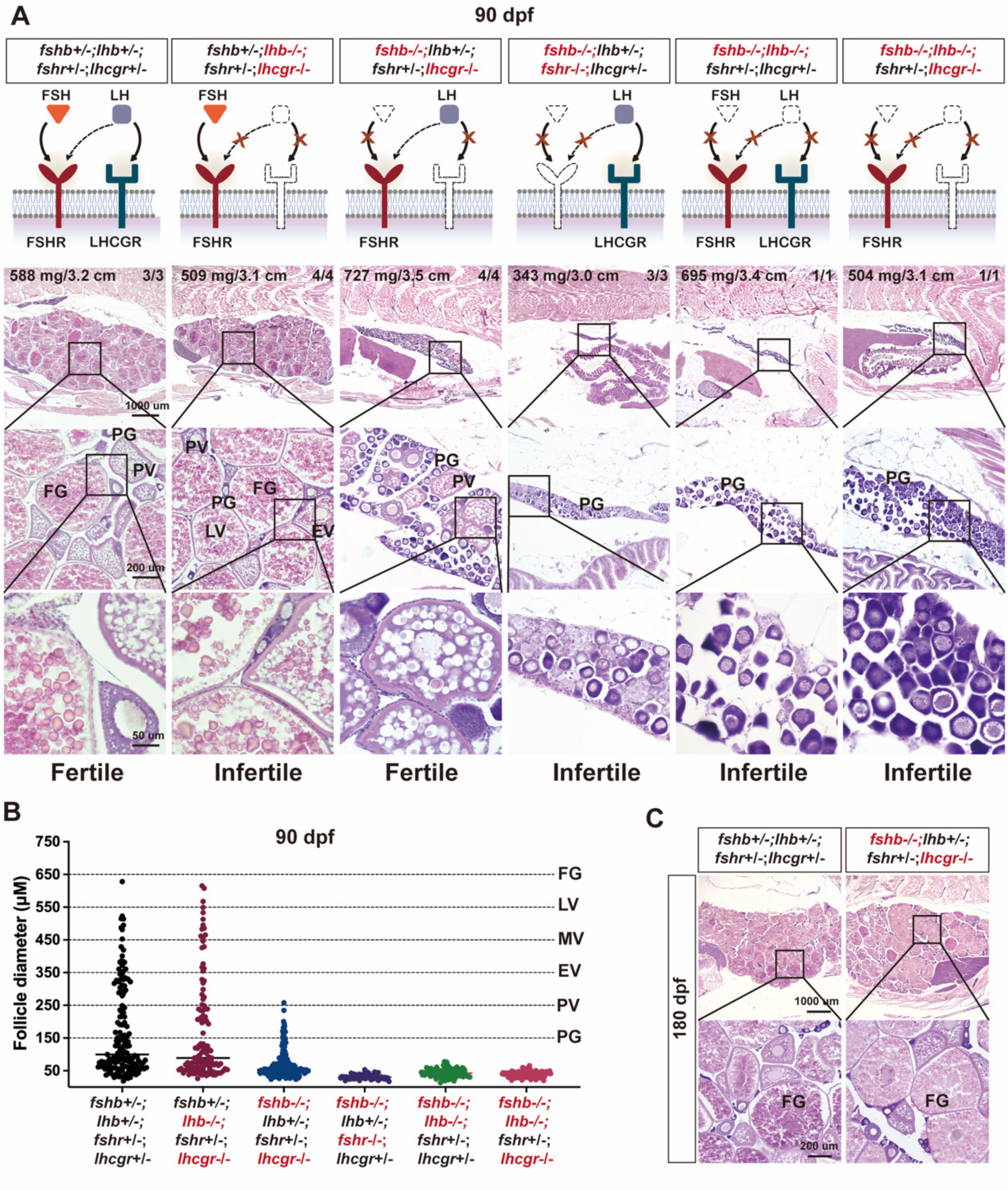
Histological examination of ovarian development and folliculogenesis in double and triple mutants. (A) Effects of disrupting gonadotropin signaling on ovarian growth and folliculogenesis at 90 dpf. Follicle stages: PG, primary growth; PV, previtellogenic stage; EV, early vitellogenic stage; MV, mid-vitellogenic stage; LV, late vitellogenic stage; FG, full-grown stage. (B) Distribution of follicle stages across genotypes at 90 dpf. The follicles in *lhb-/-;lhcgr-/-* mutants could grow to FG stage (∼650 μm), while *fshb-/-;lhcgr-/-* follicles only reached PV stage (∼250 μm). The *fshb-/-;fshr-/-*, *fshb-/;lhb-/-*, and *fshb-/-;lhb-/-;lhcgr-/-* mutant follicles were all arrested at PG stage (<150 μm). (C) Histological analysis of *fshb-/-;lhcgr-/-* ovaries at 180 dpf.

It is worth noting that we discovered one female fish in 16 double ligand mutants (1/16) (*fshb*-/-*;lhb*-/-) and one in 9 triple mutants (*fshb*-/-*;lhb*-/-*;lhcgr-/-*) (1/9), different from our previous conclusion that double ligand mutation of *fshb* and *lhb* resulted in an all-male phenotype [35], suggesting incomplete penetrance of the genotype. Histological analysis of these females showed an extreme ovarian hypotrophy with follicles completely blocked at early PG stage (Fig. 5A-B), indicating that although the two receptors (Fshr and Lhcgr) could drive spermatogenesis in the absence of ligands, their spontaneous activities were not sufficient to drive ovarian differentiation and folliculogenesis.

### Triple mutants unveil ligand-independent activity of individual receptors

Phenotype analysis of double mutants supports the notion that two gonadotropin receptors (Fshr and Lhcgr) have spontaneous or constitutive ligand-independent activities. However, such activities of individual receptors remain to be further elucidated. To address this issue, we generated all four triple mutants in zebrafish. First, all triple mutants developed as males except one female in *fshr* only fish (1/9, 11.1%) (Fig. 7). Histological analysis at both 90 dpf (Fig. 6) and 360 dpf (Fig. S3) showed that all of them exhibited significantly reduced testis size compared with the control fish (Fig. 3A). In the absence of two receptors and one ligand (either FSH or LH), the testes were extremely underdeveloped with no or limited meiotic activity, comprising primarily spermatogonia and limited number of spermatocytes with very few mature spermatozoa (Fig. 6A-C, Fig. 3B and Fig. S4). As the result, both triple mutants were infertile, failing to induce spawning of WT females (Fig. 3C). In contrast, the triple mutants retaining a single receptor, Fshr (*fshb*-/-*;lhb*-/-*;fshr*+/-*;lhcgr*-/-) or Lhcgr (*fshb*-/-*;lhb*-/-*;fshr*-/-*;lhcgr*+/-), both exhibited a significantly enhanced spermatogenic activity compared to those without any receptors (Fig. 6D-E), except a few individuals showing complete lack of meiotic activity at both 90 and 360 dpf (Fig. S4). Although the testis sizes of these mutants were similar to those of other triple mutants, both meiotic activity and production of mature sperm increased significantly (Fig. 3B). Therefore, they were partially fertile (Fig. 3C). The testis size and production of mature sperm continued to increase at 360 dpf, especially in the presence of Fshr (Fig. 3D-E).

**Fig 6.**
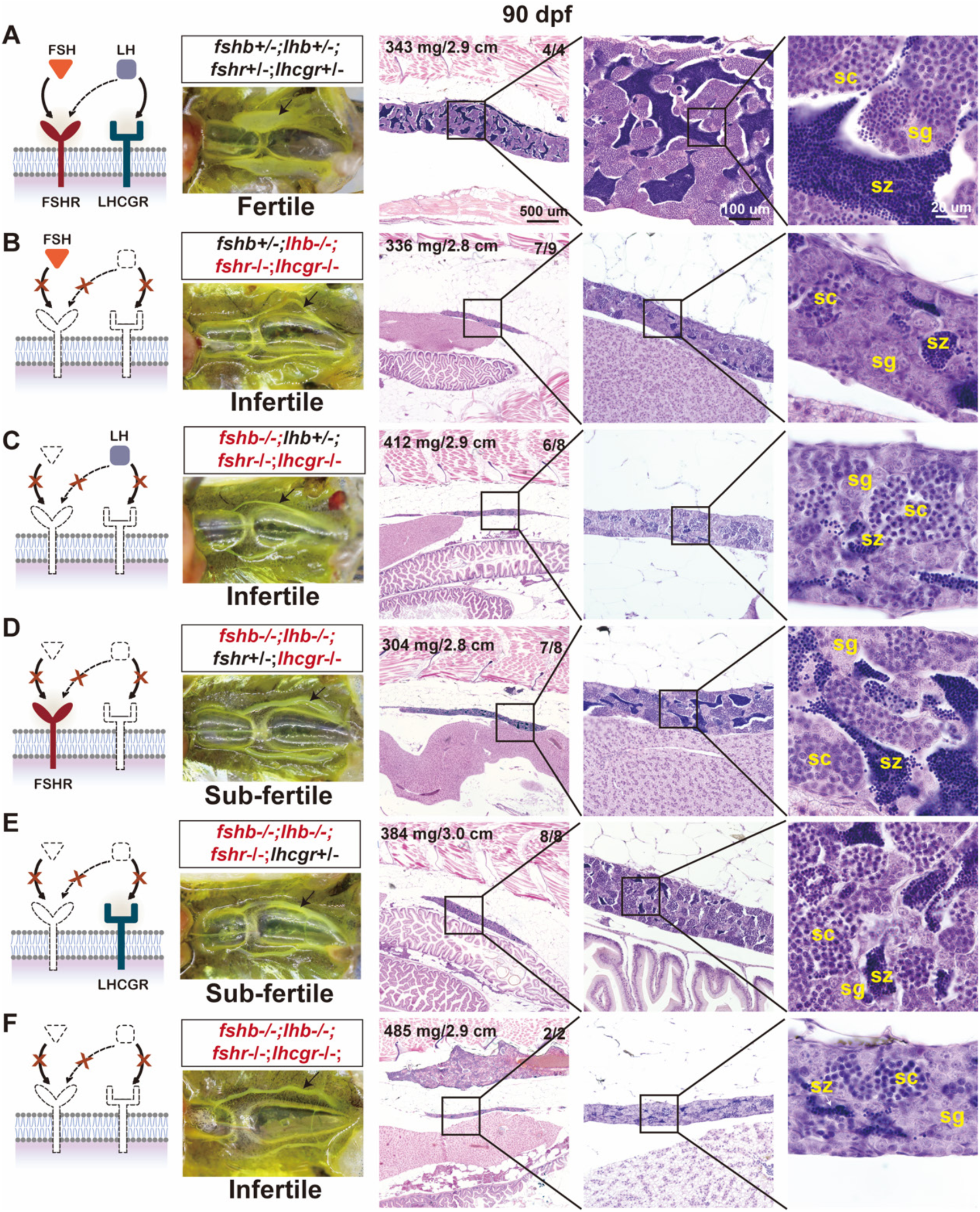
Comparative histological analysis of triple and quadruple mutant testes at 90 dpf. (A) Control testes with normal spermatogenesis. (B, C) Triple mutants with one ligand only. (D, E) Triple mutants with one receptor only. (F) Quadruple mutants with complete loss of functional gonadotropin signaling. Black arrows indicate the testes in the abdominal cavity. The body weight (mg) and body length (cm) of the representative fish are shown at the upper-left corner of the HE-stained image, while the numbers at the upper-right corner indicate the number of fish examined (lower) and those showing the same or similar phenotype (upper). Fish with discrepant divergent phenotypes are shown in Fig. S4. sg, spermatogonia; sc, spermatocytes; sz, spermatozoa.

**Fig 7.**
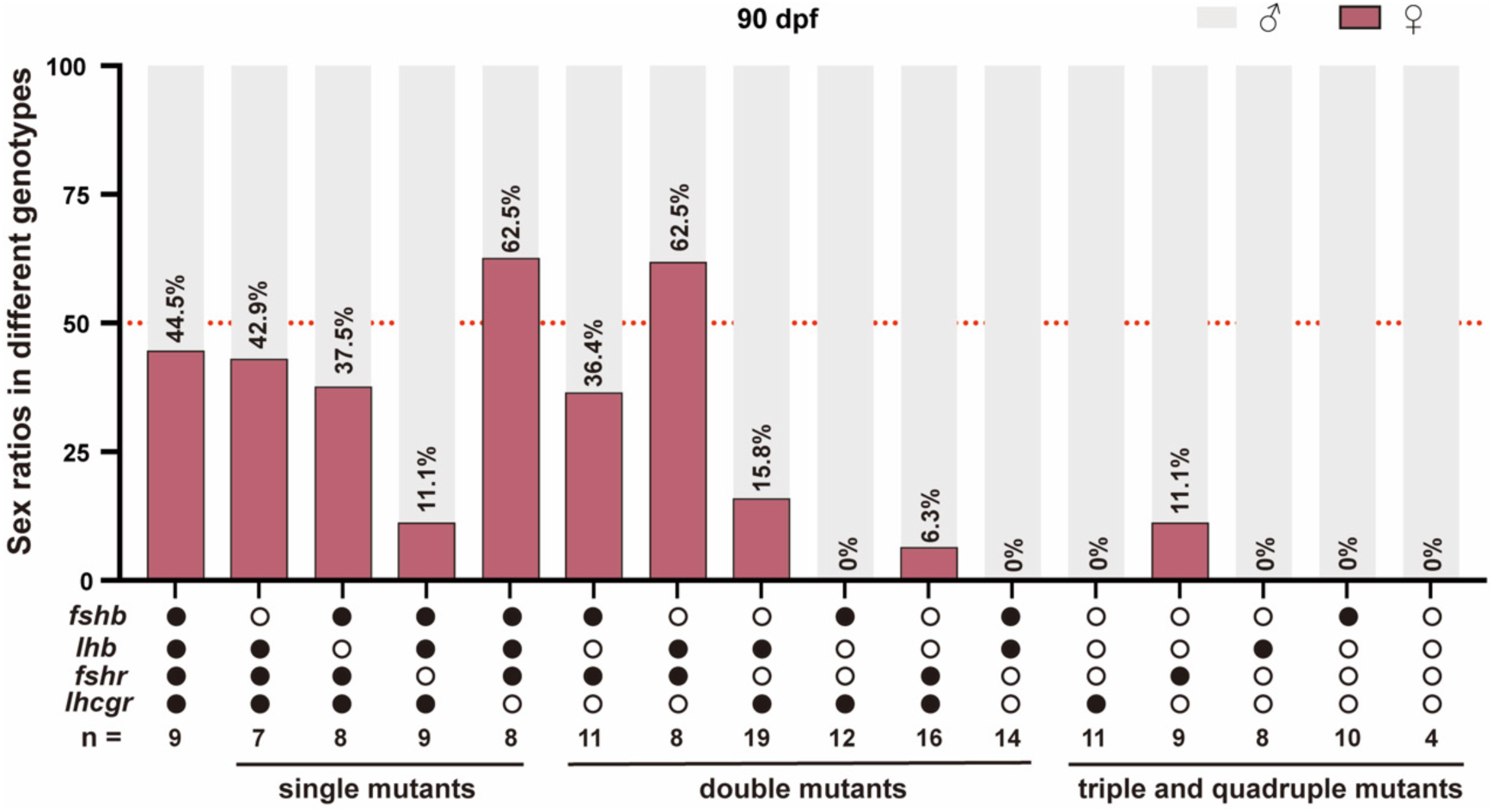
Evidence for gonadotropin involvement in sex differentiation. Sex ratios of the single, double, triple and quadruple mutants for gonadotropins (*fshb* and *lhb*) and their receptors (*fshr* and *lhcgr*) at 90 dpf. The total number of fish examined was 164, and numbers of different genotypes are shown at the bottom. ○: -/-; ●: +/-.

### Quadruple mutant highlights indispensable roles of gonadotropins in reproduction

To demonstrate the systemic impact of ablating the entire gonadotropin signaling on zebrafish reproduction, we generated a quadruple mutant with all gonadotropin ligands and receptors removed (*fshb-/-;lhb-/-;fshr-/-;lhcgr-/-*). As expected, these mutant fish exhibited normal somatic growth, with body length and weight comparable to controls and other mutant combinations at both 90 dpf and 360 dpf (Fig. S5). Histological analysis at both 90 and 360 dpf revealed that the quadruple mutant developed as all males; however, their testes were extremely underdeveloped with abundant spermatogonia and no or limited meiotic activity (Fig. 6F, Fig. S3 and Fig. S4). Consequently, the quadruple mutant males were entirely infertile, incapable of inducing spawning of WT females (Fig. 3C). It is worth noting that the phenotypes exhibited by the quadruple mutant were generally comparable to those of double mutants lacking two receptors and triple mutants with only one ligand (Fig. S2G and Fig. S3D-F).

### Genetic evidence for roles of gonadotropin signaling in sex differentiation

Gonadotropins are generally believed to play fundamental roles in regulating gonadal growth and gametogenesis during and after sex maturation in both males and females. Our results in this study also provided strong genetic evidence for roles of gonadotropin signaling in influencing sex or gonadal differentiation, especially the signaling via Fshr. In *fshr-/-* single mutant, the female ratio dropped dramatically to 11.1% as compared to 44.5%, 42.9%, 37.5%, and 62.5% for control (+/- for all four genes), *fshb-/-, lhb-/-,* and *lhcgr-/-* single mutants, respectively. The importance of Fshr in controlling sex ratio was further supported by observations in other mutant combinations (double, triple and quadruple) that whenever *fshr* was absent, the female ratio was either very low or zero. In contrast, *lhcgr* did not appear to have obvious effect on sex ratio (Fig. 7).

Analysis of double mutants also supported importance of gonadotropin signaling in sex differentiation. The lack of the two gonadotropin receptors resulted in an all-male phenotype. By comparison, in the double mutant of ligands, females showed up despite low percentage (6.3%), which was likely due to the spontaneous activities of the receptors. Whenever one ligand and one receptor were present except FSH+Lhcgr, the female ratio increased significantly (LH+Lhcgr 15.8%, FSH+Fshr 36.4%, and LH+Fshr 62.5%). It was obvious that Fshr played an important role in promoting female development. It is worth noting that FSH and Lhcgr together produced no females, again supporting our view that FSH does not activate Lhcgr (Fig. 7).

### Transcriptome analysis for differential roles of FSH and LH in oocyte maturation

One of the most striking and intriguing discoveries from the present study was that the double mutant females with the non-canonical pathway LH-Fshr (*fshb-/-;lhcgr-/-*) could undergo normal oocyte maturation and ovulation, therefore being fertile; however, the mutant females with the canonical pathway FSH-Fshr (*lhb-/-;lhcgr-/-*) were infertile although both involved signaling through Fshr. This finding raises an intriguing question about how the two pathways (FSH-Fshr, LH-Fshr), distinctly regulate oocyte maturation and ovulation via the same receptor Fshr. To address this, we performed a transcriptome analysis on FG follicles obtained from the two double mutants at 180 dpf: *lhb-/-;lhcgr-/-* (infertile due to lack of oocyte maturation) and *fshb-/-;lhcgr-/-* (fertile) (Fig. 8). The samples were collected when the follicles had started maturation but prior to germinal vesicle breakdown (GVBD) and ovulation. Before sample processing and sequencing, we performed histological and microscopic analyses to ensure healthy and comparable ovarian morphology across genotypes, with FG follicles maintaining normal shape and size (>650 μm) in both control and mutants (Fig. 8A). We also tested fecundity of the two double mutants (*lhb*-/-*;lhcgr*-/- and *fshb*-/-*;lhcgr*-/-) before sampling, and results showed infertility for *lhb*-/-;*lhcgr*-/- and reduced fertility for *fshb*-/-;*lhcgr*-/- (Fig. 8B).

**Fig 8.**
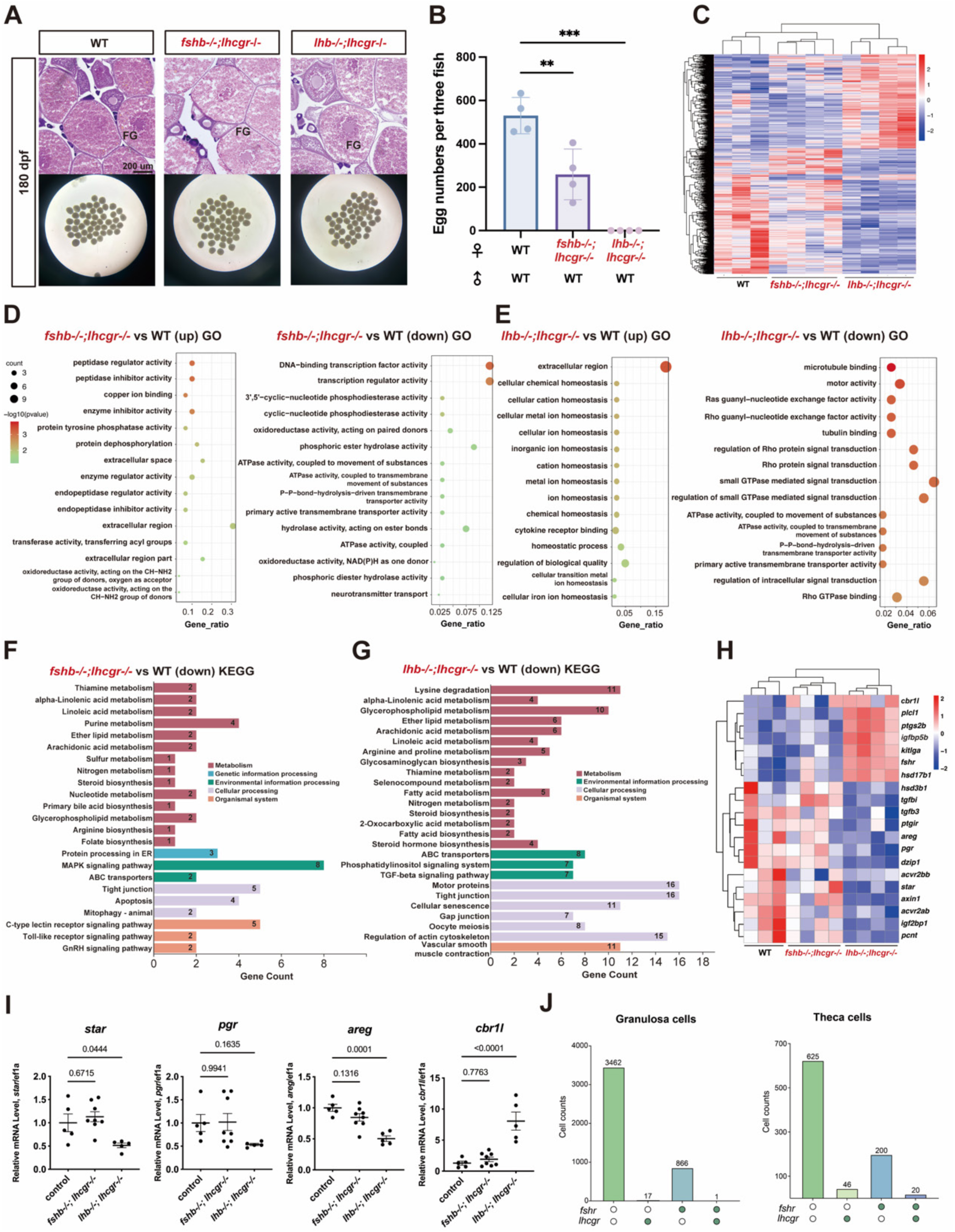
Transcriptome analysis of double mutant follicles from *lhb-/-;lhcgr-/-* and *fshb-/-;lhcgr-/-* females. (A) Morphological and histological verification of FG follicles sampled. (B) Fecundity tests at 180 dpf. Values are presented as means ± SEM and analyzed by one-way ANOVA (** *p* < 0.01 and *** *p* < 0.001). (C) Heatmap of DEGs identified. (D, E) Top enriched GO terms for all DEGs. (F, G) Top enriched KEGG pathway for down-regulated DEGs. (H) Heatmap of key DEGs with potential relevance to oocyte maturation and ovulation. (I) qPCR validation of selected genes. (J) Number of granulosa and theca cells expressing *fshr,* or *lhcgr,* or both (*fshr* and *lhcgr*) [49].

The RNA-seq data revealed a total of 532 up-regulated and 815 down-regulated genes in *lhb*-/-*;lhcgr*-/- (FSH-Fshr) FG follicles, and a total of 77 up-regulated and 114 down-regulated genes in *fshb*-/-*;lhcgr*-/- (LH-Fshr) FG follicles compared with the FG follicles from the control fish. The hierarchical clustering showed that the *fshb*-/-*;lhcgr*-/- FG follicles clustered closer to the control, whereas the *lhb*-/-*;lhcgr*-/- follicles formed a distinct group (Fig. 8C). Further analysis of DEGs showed little change GO terms between the fertile *fshb*-/-*;lhcgr*-/- (LH-Fshr) and control follicles (Fig. 8D); however, in the infertile *lhb*-/-*;lhcgr*-/- follicles, we observed profound downregulation of pathways essential for signal transduction, including small GTPase mediated signal transduction, Ras guanyl-nucleotide exchange factor activity, and Rho GTPase binding (Fig. 8E). In the *fshb-/-;lhcgr-/-* group, the MAPK signaling pathway exhibited a greater number of significantly altered genes compared with other signaling pathways (Fig. 8F). Pathway analysis linked *lhb*-/-*;lhcgr*-/- to reduced cellular processes that are potentially important for oocyte maturation, including steroid biosynthesis, oocyte meiosis, phosphatidylinositol signaling system, and TGF-β signaling pathway (Fig. 8G). We further analyzed some specific genes involved in gonadal development, follicle maturation and ovulation. The results are visualized using a heatmap, highlighting the genes whose expressions were significantly higher in the control and *fshb*-/-*;lhcgr*-/-(LH-Fshr, fertile) follicles but lower in *lhb*-/-*;lhcgr*-/- (FSH-Fshr, infertile) follicles (*star, hsd3b1, pgr, areg, acvr2ab, acvr2bb,* and *tgfb3*) and those showing the opposite changes (*ptgs2b, igfbp5b, hsd17b1, kitlga,* and *fshr*) (Fig. 8H). Validation of selected genes by qPCR confirmed the results of the transcriptome data. The expressions of *star, pgr,* and *areg* were significantly decreased, whereas *cbr1l* was upregulated in *lhb*-/-*;lhcgr*-/- follicles (Fig. 8I).

Given the functional importance of Fshr as compared to Lhcgr in zebrafish gonadal development and maturation, we also analyzed the distribution of the two receptors in follicle cells, namely granulosa and theca cells, using the data from a published single-cell transcriptomic database on 40-dpf follicles [49]. The results revealed that *fshr* was expressed in both granulosa and theca cells, whereas the expression of *lhcgr* was primarily limited to theca cells. The abundance of cells expressing *fshr* was much higher than that expressing *lhcgr,* probably due to the early stage of follicles used for the analysis. About 30% of the *lhcgr*-expressing theca cells also expressed *fshr,* which accounted for about 9% of *fshr*-expressing theca cells (Fig. 8J).

## Discussion

By systematically analyzing the phenotypes of all possible mutant combinations of gonadotropins (*fshb* and *lhb*) and their receptors (*fshr* and *lhcgr*) including single, double, triple, and quadruple mutants, this study provided valuable insights into the functional landscape of gonadotropins in controlling zebrafish reproduction in both females and males, including ligand-receptor specificity, ligand-dependent and independent receptor activities, functional roles and significance of each ligand-receptor pathway with particular reference to sex differentiation, gametogenesis, and final oocyte maturation. Our research represents the most extensive genetic analysis to date of gonadotropins and their signaling in vertebrates, providing valuable insights into both conserved and species-specific mechanisms governing reproductive regulation.

### Ligand-receptor specificity and pathway analysis

In mammals, gonadotropins and their cognate receptors interact with high specificity (FSH-FSHR and LH-LHCGR), with no cross-reactivity occurring under physiological conditions [2, 50]. In contrast, teleosts often exhibit promiscuous gonadotropin-receptor interactions, challenging the traditional “one ligand-one receptor” paradigm; however, the receptor specificity in most species remains poorly characterized. Using recombinant gonadotropins (FSH and LH) and their receptors (Fshr and Lhcgr), we previously demonstrated that zebrafish LH was able to activate both Fshr and Lhcgr, whereas FSH acted exclusively on Fshr in vitro. This finding suggests the existence of three potential signaling pathways in zebrafish: two canonical pathways (FSH-Fshr and LH-Lhcgr) and one non-canonical pathway (LH-Fshr). However, the biological relevance and functional significance of these ligand-receptor pathways have not been well understood in vivo. To address this, we employed a gene-editing approach (TALEN) to generate mutants for both gonadotropins (*fshb*-/- and *lhb*-/-) and their receptors (*fshr*-/- and *lhcgr*-/-). Phenotypic analyses of these mutants, including double ligand mutants and double receptor mutants, provided critical insights into the roles of individual gonadotropins and their receptors in controlling reproductive functions in both males and females [35, 36]. These findings are further supported by studies from other researchers [37, 51, 52].

In this study, we generated all six possible double mutants to investigate the functionality of each potential gonadotropin signaling pathway. In addition to the double ligand mutant and double receptor mutant we reported previously [35, 36], we created four additional double mutants, each with one ligand and one receptor deleted, leaving only a single pathway intact. This approach allowed us to systematically examine ligand-receptor specificity and the role of each individual pathway in regulating male and female reproduction, including the three potential pathways we proposed (FSH-Fshr, LH-Lhcgr, and LH-Fshr) and one hypothetical pathway that our previous study did not support (FSH-Lhcgr) [33, 34].

In the presence of FSH-Fshr pathway alone, both ovary and testis could grow normally with normal folliculogenesis and spermatogenesis, indicating the importance of this pathway in supporting gonadal growth in both sexes. However, the females were infertile without oocyte maturation and ovulation due to lack of LH. In contrast, the LH-Lhcgr pathway was not able to support follicle development in the ovary although it could stimulate spermatogenesis in the testis. All follicles in the ovary were arrested at early PG stage without activation and vitellogenic growth, and females were therefore also infertile. Interestingly, the non-canonical pathway LH-Fshr was the only one that could support both gonadal growth and maturation in males and females despite a delayed follicle growth and slightly reduced testis size and sperm production. This strongly supports the notion we proposed previously that LH can activate Fshr to some extent in addition to its cognate receptor Lhcgr [34].

As for the FSH-Lhcgr pathway, our previous study with recombinant hormones and receptors did not support the existence of such a pathway in zebrafish [33, 34]. However, a subsequent study suggested that FSH could also cross-activate Lhcgr to support male reproduction, evidenced by normal spermatogenesis and fertility in *lhb-/-;fshr-/-*zebrafish [51]. In contrast, genetic evidence from the present study supports our previous findings in vitro. The double mutant *lhb-/-;fshr-/-* males exhibited significantly smaller testes and reduced fertility due to impaired spermatogenesis, closely resembling the phenotypes of double ligand mutant *lhb-/-;fshb-/-* and double receptor mutant *lhcgr-/-;fshr-/-*. Furthermore, the similar levels of spermatogenesis and fertility observed in *lhb-/-;fshr-/-* (with FSH-Lhcgr) and *fshb-/-;lhb-/-;fshr-/-* (with only Lhcgr) triple mutants indicate no enhancement of Lhcgr activity by the presence of FSH, further excluding any functional relevance of FSH-Lhcgr interaction in vivo. The reported spermatogenesis and fertility in *lhb-/-;fshr-/-* zebrafish [51] were likely due to the ligand-independent spontaneous receptor activity of Lhcgr, which we also demonstrated in this study (see discussions below).

Overall, our findings confirm the existence of three functional signaling pathways: the canonical FSH-Fshr and LH-Lhcgr pathways, and the non-canonical LH-Fshr pathway. Importantly, the non-canonical LH-Fshr signaling is sufficient to support both gonadal growth and maturation in both sexes. This promiscuous receptor recognition likely reflects an evolutionary adaptation for poikilothermal animals to dynamic environmental conditions, such as temperature, salinity, and oxygen, which may affect hormone levels. This adaptation may represent a strategy that allows the same gonadotropin to perform multiple roles, thereby increasing reproductive success. This promiscuity might also be a remnant of ancestral reproductive systems, where fewer specialized hormones and receptors existed. It provides redundancy, ensuring that critical reproductive processes like gametogenesis and steroidogenesis can proceed even if one gonadotropin or receptor is impaired.

### Spontaneous constitutive receptor activities

Constitutive activity refers to the capacity of a receptor to produce biological responses in the absence of ligands. This phenomenon is observed in various receptors, including G-protein-coupled receptors [53, 54], and is mechanistically explained by the two-state model, wherein receptors spontaneously shift to an active state without ligand binding. For instance, constitutive activity of FSHR has been reported in mice with a role of maintaining normal Leydig cell development in FSHβ-deficient mice [40]. Similarly, Lhcgr in channel catfish has also been suggested to exhibit ligand-independent activity in vitro [42]. In our previous study, we also observed spontaneous receptor activity for zebrafish Lhcgr in vitro [33].

In this study, we offered solid genetic evidence for the ligand-independent activity of both Fshr and Lhcgr in vivo. The double ligand mutant *fshb*-/-*;lhb*-/- (with two receptor) and triple mutants *fshb*-/-*;lhb*-/-*;fshr*-/- (with only Lhcgr) and *fshb*-/-*;lhb*-/-*;lhcgr*-/-(with only Fshr) all exhibited significant spermatogenic activities, leading to production of significant amounts of mature sperm. These male fish were sub-fertile at 90 dpf compared to receptor-null double, triple and quadruple mutants, which were completely infertile. The impact of constitutive activity became more unequivocal at later stage (360 dpf): the males with either Fshr or Lhcgr developed even larger testes with abundant mature sperm. This phenotype was particularly pronounced when Fshr was available. Our discoveries demonstrated that Fshr or Lhcgr alone was sufficient to drive the entire process of spermatogenesis due to their spontaneous ligand-independent activities and highlighted the Fshr-centric signaling in zebrafish.

### Gonadotropin participation in gonadal differentiation

Sex determination and differentiation in zebrafish remain an intriguing issue. Unlike mammals, zebrafish does not possess heteromorphic sex chromosomes [55], suggesting different sex determination mechanisms in this species. No sex-linked markers, such as *Sry* in mammals or *dmy* in medaka (*Oryzias latipes*), have been identified in the zebrafish [56, 57]. It is generally accepted that zebrafish adopts a polygenic mechanism for its sex determination [58]. In addition to genetic factors, various environmental conditions including hypoxia [59], population density [60], temperature [61], and food availability [62] have been suggested to affect sex development in zebrafish. Besides, the number of germ cells has also been implicated in female differentiation in zebrafish [63]. However, few studies have suggested roles for gonadotropin signaling in gonadal differentiation in fish. In a hermaphroditic gobiid fish (*Trimma okinawae*), which can change sex back and forth depending on social status in the community, the expression of gonadotropin receptors is closely associated with the developmental state of the gonads during sex reversal [64]. In Japanese flounder (*Paralichthys olivaceus*), a species with sex differentiation sensitive to environmental temperature, *fshr* is only expressed in ovarian tissue during gonadal differentiation. Elevated temperatures promote testis development during gonadal differentiation, in part by suppressing *fshr* expression [65]. Although these studies suggest a role for gonadotropin signaling in gonadal differentiation, a definitive cause-effect relationship has not yet been established.

The present study provided strong genetic evidence for roles of gonadotropin signaling in gonadal differentiation, either primary or secondary differentiation (sex reversal), particularly the signaling pathways involving Fshr. Whenever Fshr was present with either FSH or LH or both, the female ratios were always comparable to that of the control (∼50% with all ligands and receptors present). The female ratio dropped precipitously in all combinations lacking Fshr, often to zero (quadruple mutant and three triple mutants without *fshr*). In contrast, the presence of Fshr alone was able to support female development with its spontaneous activity, despite low percentage (11.1%). The percentage of females increased drastically when Fshr was present with either FSH (FSH-Fshr, 36.4%) or LH (LH-Fshr, 62.5%), further supporting the notion that Fshr can be activated by both FSH and LH [34]. By comparison, Lhcgr also participated in female differentiation, but its activity was much weaker compared to Fshr. The presence of Lhcgr alone was not able to support female development, unlike Fshr; however, females appeared in the double mutant with both Lhcgr and LH (Lhcgr+Lhb), but not FSH, further supporting our previous discovery that Lhcgr could be activated by LH, but not FSH [34]. It should be noted that the lack or low percentage of females in all mutant combinations lacking *fshr* could also be due to sex reversal of differentiated females to males as we reported previously [36]. However, in quadruple mutant and three triple mutants without *fshr,* we did not observe formation of ovaries with distinct follicles, indicating a complete lack of female differentiation without sufficient gonadotropin signaling.

### Roles of gonadotropin signaling in gametogenesis

Previous studies have demonstrated essential roles of gonadotropins and their receptors in gametogenesis [35–37]. Genetic studies in different fish model suggest a relatively conserved function for FSH between fish and mammals. In zebrafish, *fshb* mutation delays folliculogenesis including both follicle activation or PG-PV transition and vitellogenic growth; however, its function can be compensated by LH through cross reaction with Fshr [35]. In medaka, loss of *fshb* results in arrested follicle development at PG stage, characterized by lack of cortical alveoli and yolk granules in oocytes [66]; it is not yet clear if LH can rescue this phenotype at later stages as observed in zebrafish although it has been reported that LH can also activate Fshr in medaka [24]. A similar study in short-lived turquoise killifish (*Nothobranchius furzeri*) demonstrated that disruption of *fshb* blocks follicle development at PV stage with formation of cortical alveoli but not yolk granules [67]. The phenotypes seen in these fish *fshb* mutants are generally consistent with those found in *Fshb* mutant mice, where follicles are blocked at the antral stage [9, 35, 66, 67].

In contrast to *fshb* mutation, mutation of *lhb* has no impact on follicle growth in both zebrafish and medaka. However, females without LH cannot undergo oocyte maturation and are therefore anovulatory in both species [35, 48, 66]. This phenotype could be rescued in medaka by either homologous recombinant medaka LH or heterologous pregnant mare serum gonadotropin (PMSG) [66] and in zebrafish by in vivo administration of human chorionic gonadotropin (hCG) and in vitro treatment of isolated follicles with the maturation-inducing hormone 17α,20β-dihydroxy-4-pregnen-3-one (DHP) [48]. This phenotype contrasts sharply with that observed in mice whose *lhb* mutation caused severe abnormalities in antral follicles, resulting in lack of healthy preovulatory follicles and corpora lutea [14].

We further demonstrated that LH-stimulated follicle growth occurred via Fshr, not Lhcgr, as normal folliculogenesis was observed in the *fshb-/-;lhcgr-/-* mutant with LH-Fshr, though at a slower rate, but not in the *fshb-/-;fshr-/-* mutant with LH-Lhcgr. The critical role of Fshr in folliculogenesis is supported by the significant increase in *fshr* expression during the PG-PV transition and its continued upregulation in the vitellogenic growth stage [33, 44, 45]. Our findings in the present study reaffirmed the substitutability of FSH by LH, the indispensability of Fshr, and the insignificance of Lhcgr during zebrafish folliculogenesis. The essential role of Fshr in regulating folliculogenesis has also been demonstrated in medaka using loss-of-function approach. An early investigation using target-induced local lesions identified a medaka *fshr* mutant that exhibited a phenotype comparable, though not identical, to that observed in zebrafish [36]. Female medaka *fshr* mutants were infertile, with follicles arrested at the PG or early PV stage with signs of cortical alveolar formation [68], in contrast to zebrafish *fshr* mutant, which showed extremely underdeveloped ovaries with early PG follicles only [36]. Consistent with our observations in zebrafish, certain *fshr* mutant XX females in medaka also underwent sex reversal, developing male characteristics [36, 68].

The present study also provided substantial evidence for the indispensable roles of gonadotropin signaling in testis development and spermatogenesis. This is well demonstrated by double, triple, and quadruple mutants lacking both *fshr* and *lhcgr*. The males of all these mutants were infertile with extremely small testes and limited spermatogenic activity, which agrees well with previous observations in the double receptor mutant [36, 37]. Interestingly, despite the underdeveloped testes and impaired spermatogenesis, we observed the formation of mature spermatozoa in all mutant combinations without the two receptors, albeit at extremely low levels. This finding extends our previous report on the double receptor mutant, which showed extremely underdeveloped testes with limited meiotic activity [36]. It is worth noting that the presence of either Fshr or Lhcgr in triple mutants significantly increased spermatogenic activity, resulting in subfertility, further supporting the ligand-independent constitutive or spontaneous receptor activities for both Fshr and Lhcgr [33, 37]. The role and importance of Fshr in controlling testis growth and spermatogenesis have also been reported recently in Atlantic salmon. Mutation of salmon *fshr* using CRISPS/Cas9 resulted in an extremely underdeveloped testis with complete arrest of spermatogenic cells at the stage of type A spermatogonia (SGA) without signs of meiosis [69].

The presence of mature spermatozoa in the quadruple mutant male zebrafish indicates that spermatogenesis can take place independently of gonadotropin signaling, despite the testes being significantly underdeveloped and spermatogenesis severely impaired. This raises an interesting question about what other hormones or factors are involved in supporting basal spermatogenesis in the absence of gonadotropin signaling. Local growth factors may likely play critical roles in this process. An early study reported that insulin-like growth factor 1 (IGF-1) alone could promote spermatogonial differentiation into spermatocytes and spermatids in cultured zebrafish testes [70]. Another potential regulator is anti-Müllerian hormone (Amh). Our recent studies proposed that Amh might serve dual functions in zebrafish spermatogenesis: it inhibits spermatogonial proliferation by suppressing gonadotropin signaling while simultaneously promoting their differentiation into meiotic spermatogenic cells [71, 72]. Our recent study further supports a critical role for Amh in testicular meiosis and spermatogenesis. Simultaneous mutation of *amh* and gonadotropin receptors (*fshr* and *lhcgr*) completely arrested spermatogenesis at the spermatogonial stage, with no signs of meiosis [71].

### Differential roles and mechanisms of FSH and LH in final oocyte maturation

One of the most striking discoveries in this study was the distinct roles of FSH and LH in promoting final oocyte maturation through the same receptor Fshr. Analysis of double mutants demonstrated that the FSH-Fshr and LH-Fshr pathways in *lhb-/-;lhcgr-/-* and *fshb-/-;lhcgr-/-* double mutants exhibited different phenotypes regarding oocyte maturation and ovulation, despite both ligands activating the same receptor Fshr in the absence of Lhcgr. Both pathways supported ovarian growth and folliculogenesis; however, while the females with LH-Fshr could spawn normally, those with FSH-Fshr could not lay eggs and were therefore infertile, similar to the single *lhb-/-* mutant. This phenomenon where a receptor responds differently to different ligands regarding downstream signaling mechanisms and functions is known as biased agonism or ligand bias, which has been well documented in mammals, especially among G protein-coupled receptors [73, 74]. This phenomenon of biased agonism or functional selectivity has also been reported for gonadotropin receptors (FSHR and LHCGR), allowing the same receptor to respond differently to different ligands. For example, both human LH and chorionic gonadotropin (hCG) act as natural biased agonists for the same receptor LHCGR. While both are full agonists for cAMP and testosterone production, LH demonstrates significant partial agonism for β-arrestin recruitment and subsequent progesterone synthesis when compared to hCG [75]. Similarly, biased agonism also occurs at FSHR, where the glycosylation pattern of FSH influences its signaling pathways. Specifically, hypo-glycosylated FSH variants preferentially stimulate β-arrestin-dependent ERK activation, whereas fully-glycosylated forms are less active in this pathway [76]. The distinct responses of zebrafish Fshr to FSH and LH in terms of oocyte maturation provides an excellent example for biased agonism in fish. To further explore the downstream mechanisms of FSH and LH activation of Fshr, resulting in different responses in oocyte maturation, we performed a transcriptome analysis of full-grown follicles collected prior to oocyte maturation from control and the two double mutants *lhb-/-;lhcgr-/-* and *fshb-/-;lhcgr-/-* with FSH-Fshr and LH-Fshr, respectively.

Transcriptome analysis showed clearly that the expression profiles of *fshb*-/-;*lhcgr*-/-FG follicles were similar to that of the control group, while both differed significantly from that of the infertile *lhb*-/-*;lhcgr*-/- group. Among the biological processes and pathways identified by GO and KEGG analyses on DEGs, we were particularly interested in pathways associated with steroid biosynthesis, oocyte meiosis, and TGF-β signaling as well as growth factors potentially involved in controlling oocyte maturation and ovulation. Many genes of these pathways, including *star, hsd3b1, pgr, acvr2ab, acvr2bb, tgfb3,* and *areg,* were significantly down-regulated in *lhb*-/-*;lhcgr*-/-follicles.

Steroidogenesis involves specific functional proteins such as steroidogenic acute regulatory protein (Star/*star*) and a series of enzymes such as 3-β-hydroxysteroid dehydrogenase (Hsd3b1/*hsd3b1*), which transports cholesterol into mitochondria for DHP synthesis and catalyzes 3β-hydroxysteroid dehydrogenation and Δ^5^- to Δ^4^-isomerisation of the Δ^5^-steroid precursors, respectively, thus leading to the formation of all classes of steroid hormones [77]. These molecules are subject to regulation by LH in mammals [78]. Pharmacological and genetic studies have demonstrated the indispensable roles of steroids in LH-induced oocyte maturation and ovulation across teleosts. In zebrafish and medaka, the maturation-inducing hormone DHP potently induces oocyte maturation and ovulation [79–81], whereas inhibitors targeting steroidogenesis (trilostane) or progestin signaling via nuclear progestin receptor (Pgr) (RU486) effectively blocked these processes in medaka [82]. Consistent with these findings, genetic ablation of membrane progestin receptors (mPRs) or Pgr results in failed oocyte maturation and ovulation, respectively [83–87]. The down-regulation of *star, hsd3b1* and *pgr* in infertile *lhb*-/-*;lhcgr*-/- but not fertile *fshb*-/-*;lhcgr*-/- and control follicles indicates that impaired steroidogenesis, which may result in reduced production of DHP for oocyte maturation and Pgr-mediated progestin signaling for ovulation could be a mechanism for unsuccessful oocyte maturation and ovulation in FSH-Fshr females without LH. This agrees well with the reports that administration of exogenous DHP could restore female fertility in zebrafish *lhb-/-* and *star-/-* mutants [47, 48, 81]. However, DHP administration was unable to rescue the anovulatory phenotype caused by *pgr* deficiency [81, 83].

The significant downregulation of amphiregulin (Areg/*areg*) in *lhb*-/-*;lhcgr*-/- mutants raises an interesting question about the involvement of epidermal growth factor (EGF) family in final oocyte maturation and ovulation. In mammals, LH-induced maturation and ovulation had remained a mystery until the discovery that LH triggered oocyte maturation and ovulation in mice via EGF family ligands, including amphiregulin (AREG), betacellulin (BTC), and epiregulin (EREG), which activated EGF receptor (EGFR) to promote cumulus expansion and oocyte maturation [88]. Subsequent work has further confirmed EGFR signaling as the central meditator that propagates signals from mural granulosa cells through cumulus cells to oocytes, activating meiotic resumption and ovulatory cascades [89]. In teleosts, our previous studies in zebrafish have provided substantial evidence for roles of EGF family and their signaling via EGFR (Egfr/*egfr*) in controlling folliculogenesis, especially oocyte maturation [90–92], which may likely involve activin system [93–95].

In addition to members of the EGF family, members of the transforming growth factor β (TGF-β) superfamily, have also been implicated in oocyte maturation in teleosts. Studies in zebrafish and killifish have demonstrated that activin promotes the acquisition of oocyte maturational competence and stimulates final maturation [96–100]. In the present study, the significant down-regulation of activin type II receptors (*acvr2ab* and *acvr2bb*) in *lhb*-/-*;lhcgr*-/- mutant follicles suggested impaired activin signaling in LH-induced oocyte maturation, which could be one of the mechanisms underlying the failure of oocyte maturation and ovulation.

Our findings from the comparative analysis of two double mutants *fshb*-/-*;lhcgr*-/- and *lhb*-/-*;lhcgr*-/- indicate that impaired LH-DHP-Pgr pathway, along with disrupted Egfr and activin signaling, in *lhb*-/-*;lhcgr*-/- mutants were likely major mechanisms that lead to failure of follicle maturation and ovulation. In contrast, the expression levels of these genes in the *fshb*-/-*;lhcgr*-/- follicles remained comparable to those in controls, allowing for proper oocyte maturation and ovulation. This highlights the specific and non-redundant role of LH in regulating the genetic pathways necessary for these processes. Interestingly, the absence of Lhcgr does not affect either follicle growth or final maturation and ovulation, further emphasizing the importance of Fshr in these pathways.

Taken together, our data strongly support a model of biased agonism at Fshr in zebrafish. We hypothesize that FSH may act as a biased agonist, primarily stimulating the canonical Gs-cAMP pathway without recruiting β-arrestin to activate the MAPK/ERK cascade. This single pathway plays a key role in promoting follicle growth but is insufficient to drive oocyte maturation. Conversely, LH, acting on the same receptor Fshr, functions as a more balanced agonist, activating both the canonical Gs/cAMP and the non-canonical G-protein-independent β-arrestin-MAPK pathways, which may also involve EGFR signaling. This dual signaling mechanism provides the complete molecular stimulus necessary to promote both oocyte growth and maturation.

## Conclusions

By generating double, triple, and quadruple mutants of gonadotropins (*fshb* and *lhb*) and their receptors (*fshr* and *lhcgr*), this study provided comprehensive evidence to clarify the roles of gonadotropin signaling in zebrafish reproductive function. The major conclusions are as follows: (1) LH cross-activating Fshr in zebrafish can compensate for the FSH role in gonadal development, with no evidence found for FSH-Lhcgr interaction; (2) Ligand-independent activity of Fshr or Lhcgr could support spermatogenesis and produce fertilizable sperm; (3) Fshr signaling is involved in female sex differentiation; and (4) Impaired LH-DHP-Pgr signaling, Egfr signaling, and activin system in the *lhb-/-;lhcgr-/-* mutants might lead to the failure of follicle maturation and ovulation. This study elucidates the complex roles of gonadotropin signaling in zebrafish reproductive biology, providing useful insights for future research.

## Acknowledgments

We thank Ms. Phoenix Un Ian LEI for the maintenance and management of the zebrafish facility and the Histology Core of the Faculty of Health Sciences for technical support. We acknowledge the Histology Core of the Faculty of Health Sciences for its invaluable technical support. This research was supported by funding from the University of Macau (MYRG-GRG2023-00144-FHS-UMDF, MYRG-GRG2024-00191-FHS, CPG2024-00030-FHS, and CPG2025-00037-FHS) and The Macau Fund for Development of Science and Technology (FDCT0086/2022/AFJ and FDCT-NSFC Joint Project 0086/2022/AFJ) to WG.

## Declaration of generative AI and AI-assisted technologies in the writing process

In preparing this work, the author(s) used Copilot to polish the language to enhance clarity and readability for certain sentences. The application of AI technology was strictly limited to language refinement and did not extend to data analysis, reasoning, referencing and conclusions. Following each use, the author(s) thoroughly reviewed and edited the content as necessary to conform to authors’ writing style and will assume full responsibility for the content of the final publication

---

**Fig S1.**
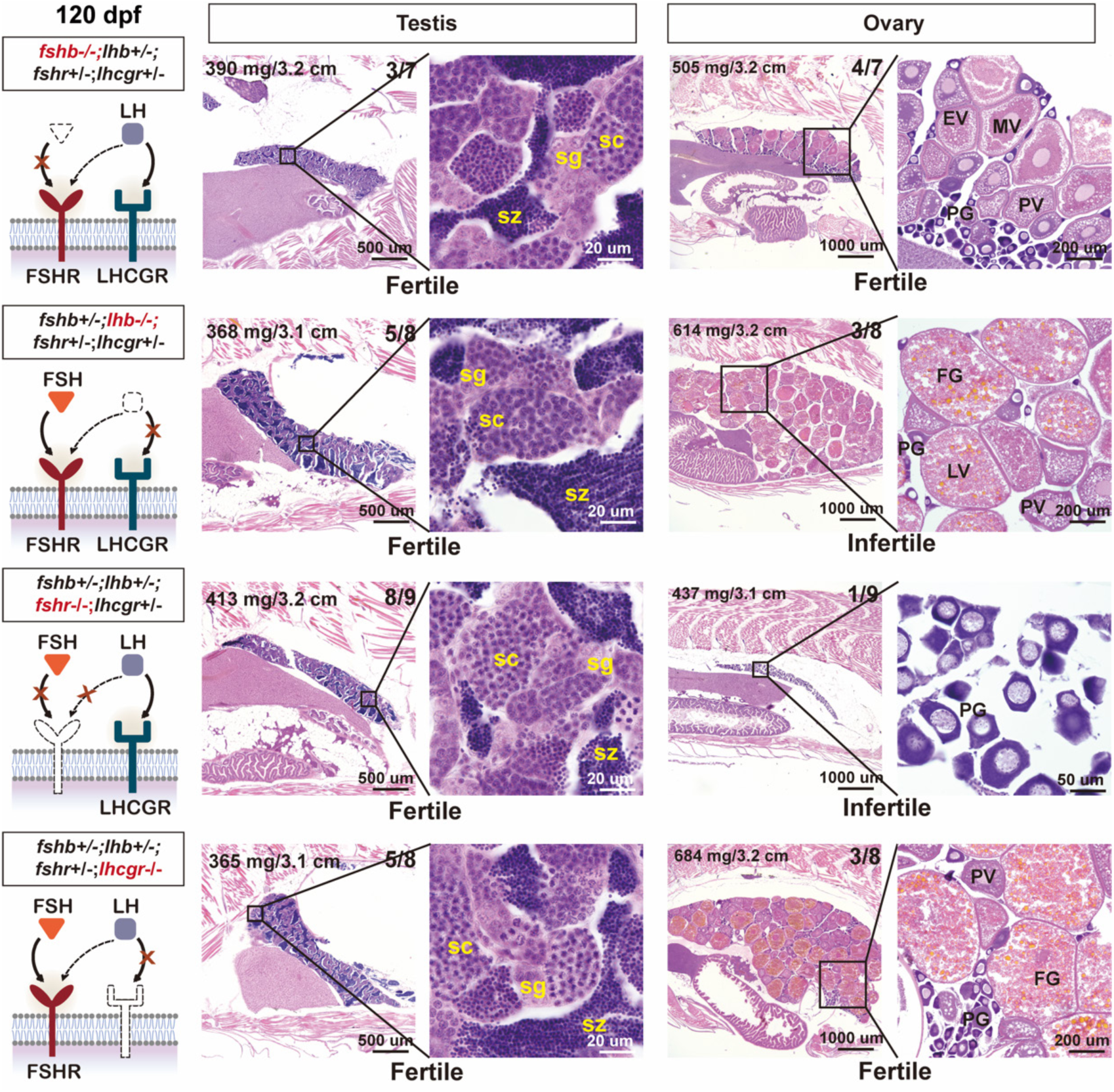
Histological analysis of single mutant males and females at 120 dpf. Absence of FSH (*fshb*) delayed ovarian and follicle growth, while loss of FSH receptor (*fshr*) completely halted follicles at the PG stage. Loss of LH (*lhb*) and its receptor (*lhcgr*) had no effect on gonadal growth and gametogenesis (spermatogenesis and folliculogenesis). sg, spermatogonia; sc, spermatocytes; sz, spermatozoa; PG, primary growth; PV, previtellogenic; EV, early vitellogenic; MV, mid-vitellogenic; LV, late vitellogenic; FG, full-grown.

**Fig S2.**
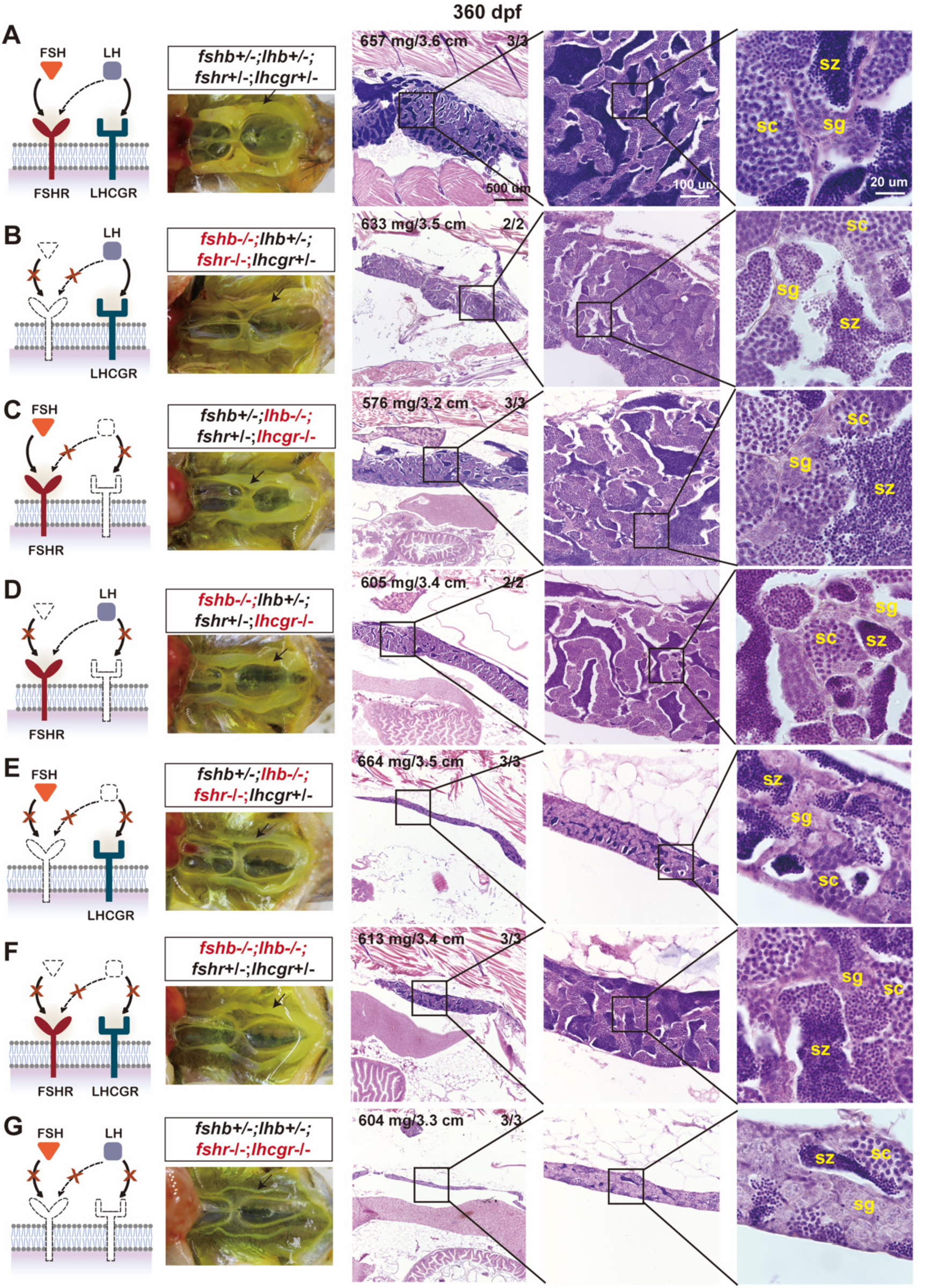
Histological analysis of testis in double mutant males at 360 dpf. (A) Control testes (*fshb+/-;lhb+/-;fshr+/-;lhcgr+/-*). (B-E) Double mutants with one pathway present. The pathways of LH-Lhcgr (*fshb-/-;fshr-/-*), FSH-Fshr (*lhb-/-;lhcgr-/-*) and LH-Fshr (*fshb-/-;lhcgr-/-*) were all sufficient to support normal testis development and spermatogenesis. However, the putative pathway FSH-Lhcgr (*lhb*-/-*;fshr-/-)* did not seem to promote testis development. (F, G) Double ligand mutants ((*fshb*-/-*;lhb*-/-) and double receptor mutants (*fshr*-/-*;lhcgr*-/-). The double ligand mutants with two receptors present showed higher spermatogenic activity than that of double receptor mutants, indicating spontaneous receptor activities. sg, spermatogonia; sc, spermatocytes; sz, spermatozoa.

**Fig S3.**
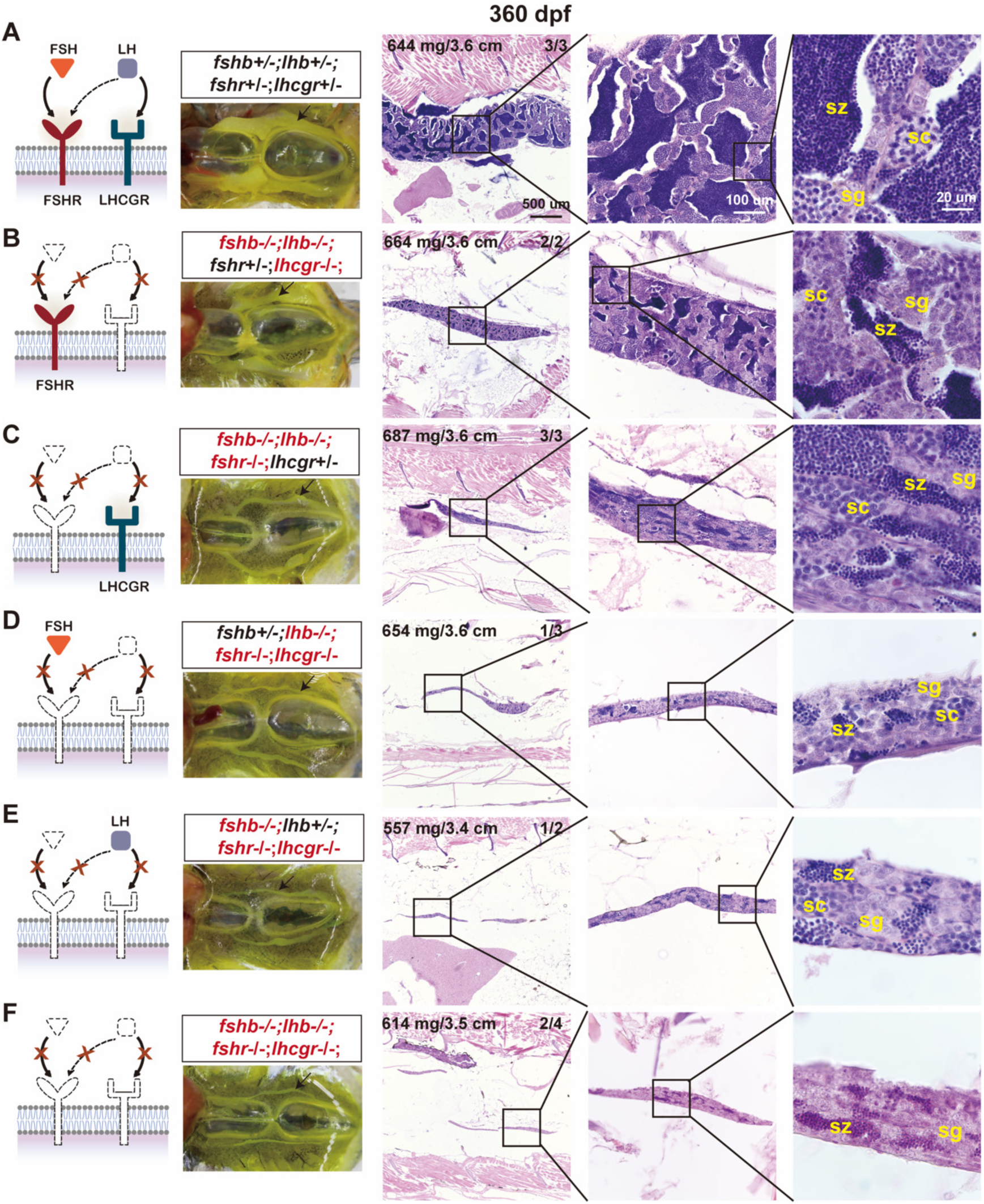
Histological analysis of triple and quadruple mutant males at 360 dpf. (A) Control testes (*fshb+/-;lhb+/-;fshr+/-;lhcgr+/-*). (B, C) Triple mutants with only one receptor. (D, E) Triple mutants with only one ligand. (F) Quadruple mutants. Fish with discrepant phenotypes are shown in Fig. S4. sg, spermatogonia; sc, spermatocytes; sz, spermatozoa.

**Fig S4.**
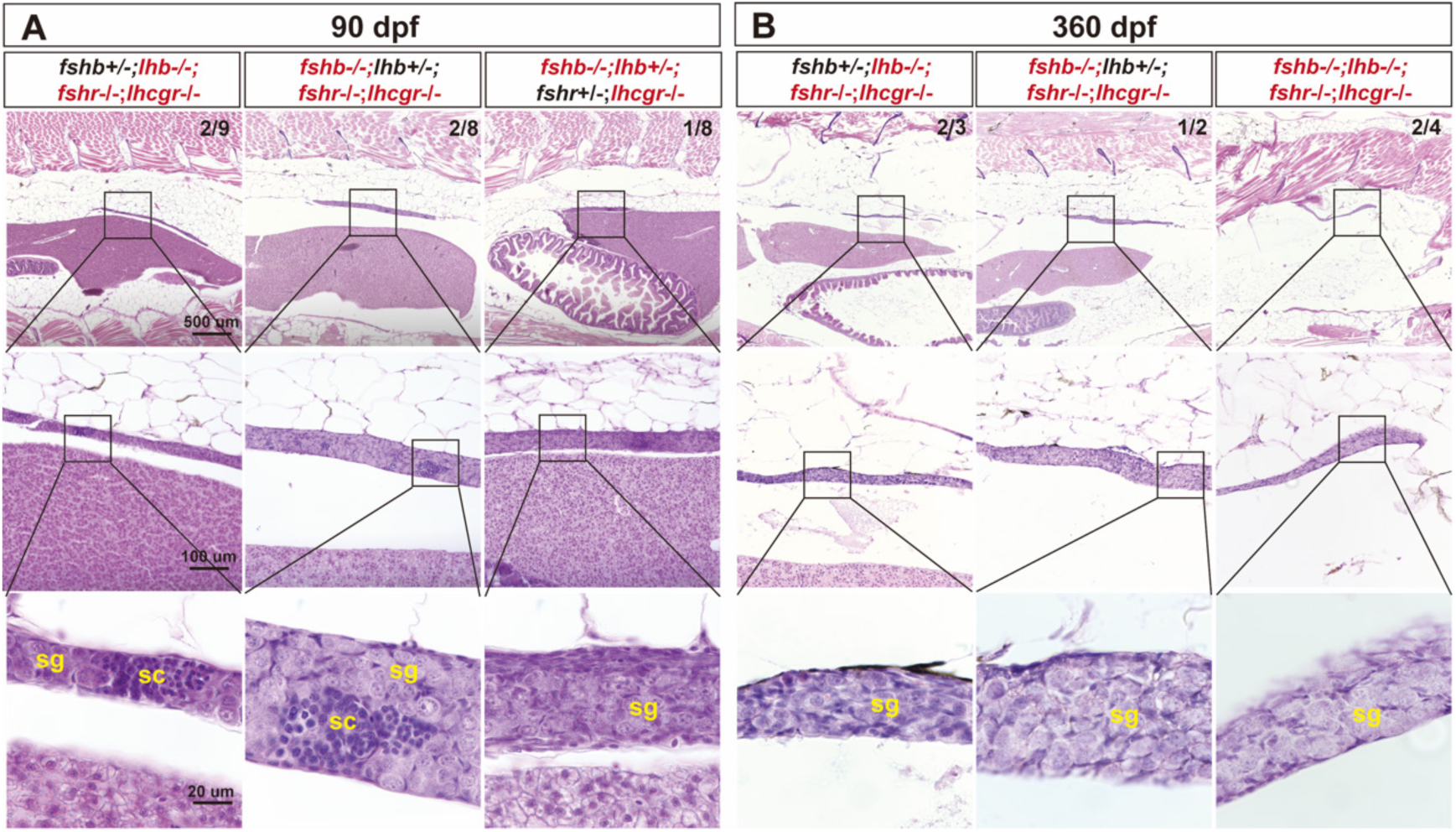
Histology of triple and quadruple mutant males with differing phenotypes. Unlike the samples in Fig. 6 and S3, some triple and quadruple mutants showed little to no meiotic activity at both 90 and 360 dpf. The upper-right numbers indicate how many fish displayed this phenotype (upper) versus the total examined (lower). sc, spermatocytes; sg, spermatogonia.

**Fig S5.**
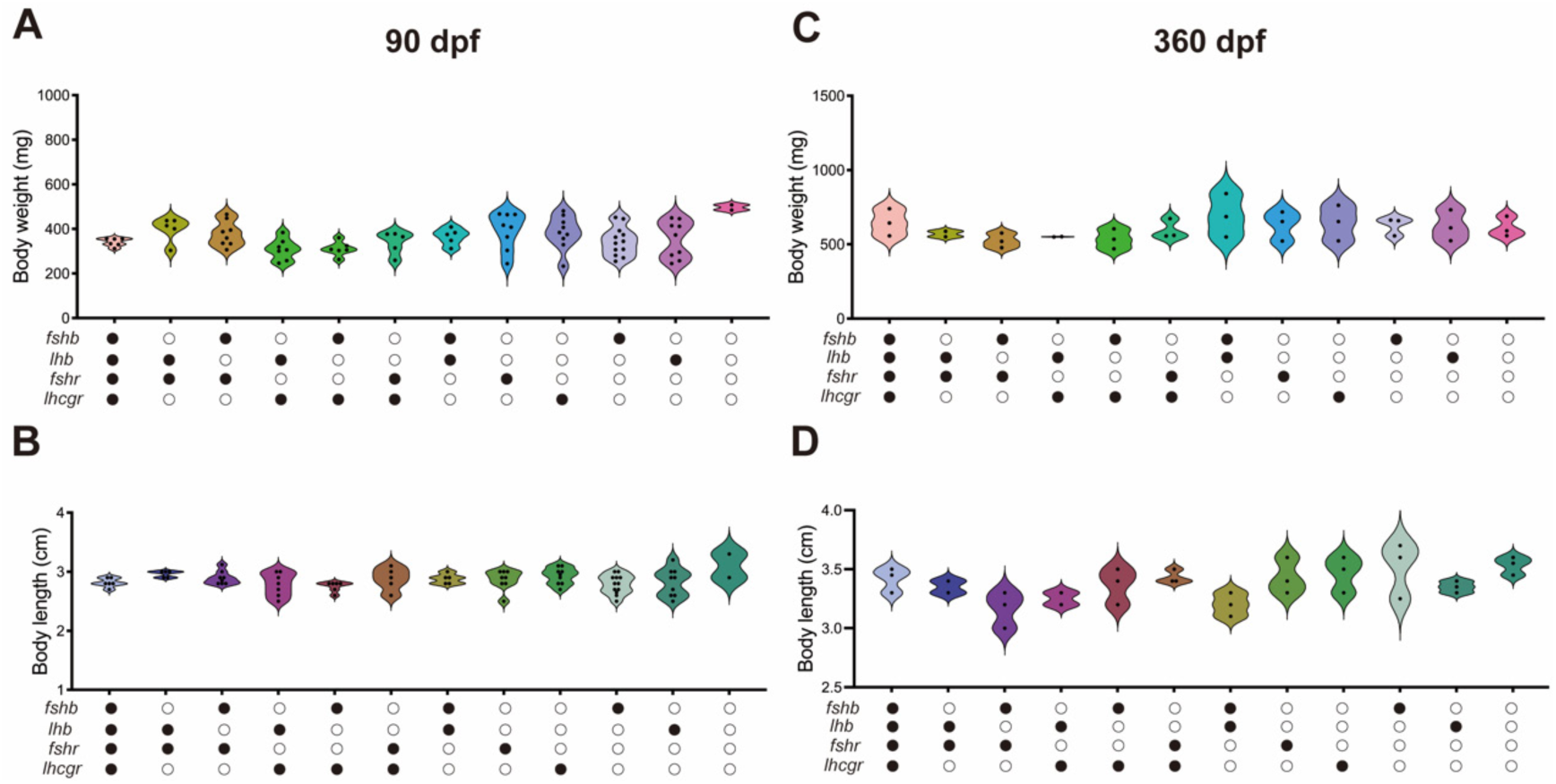
Growth performance in gonadotropin signaling-deficient mutants. (A, B) Body weight and body length at 90 dpf. (C, D) Body weight and body length at 360 dpf. Data are presented as means ± SEM and analyzed by one-way ANOVA. ○: -/-; ●: +/-.

**Table S1.**
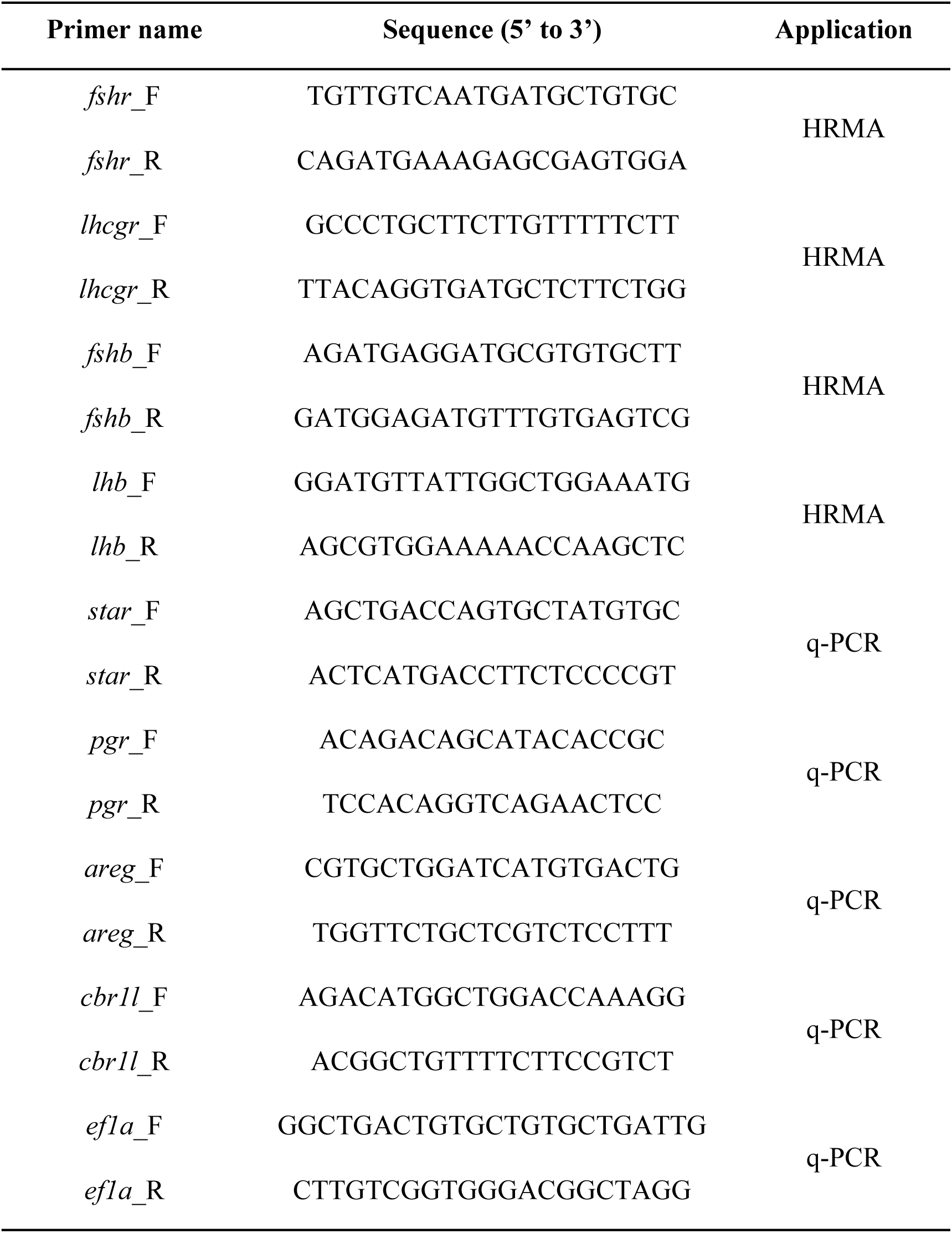
Primers used for HRMA and q-PCR.

## Notes

### Competing Interest Statement

The authors have declared no competing interest.

## References

1. Casarini L, Simoni M. Recent advances in understanding gonadotropin signaling. Fac Rev. 2021; 10:41. doi: 10.12703/r/10-41. PMID: 34046645.

2. Costagliola S, Urizar E, Mendive F, Vassart G. Specificity and promiscuity of gonadotropin receptors. Reproduction. 2005; 130:275–81. doi: 10.1530/rep.1.00662. PMID: 16123234.

3. Burns KH, Matzuk MM. Minireview: genetic models for the study of gonadotropin actions. Endocrinology. 2002; 143:2823–35. doi: 10.1210/endo.143.8.8928. PMID: 12130545.

4. Gray C. Glycoprotein gonadotropins. Structure and synthesis. Acta Endocrinol Suppl. 1988; 288:20–7. PMID: 3138864.

5. Casarini L, Crépieux P. Molecular mechanisms of action of FSH. Front Endocrinol. 2019; 10:305. doi: 10.3389/fendo.2019.00305. PMID: 31139153.

6. Azhar S, Clark MR, Menon KMJ. Regulation of cyclic adenosine 3′,5′-mono-phosphate dependent protein kinase of rat ovarian cells by luteinizing hormone and human chorionic gonadotropin. Endocr Res Commun. 1976; 3:93–104. doi: 10.3109/07435807609052925. PMID: 182451.

7. Takahashi T, Ogiwara K. cAMP signaling in ovarian physiology in teleosts: A review. Cell Signal. 2023; 101:110499. doi: 10.1016/j.cellsig.2022.110499. PMID: 36273754.

8. Wang Y, Ge W. Involvement of cyclic adenosine 3’,5’-monophosphate in the differential regulation of activin βA and βB expression by gonadotropin in the zebrafish ovarian follicle cells. Endocrinology. 2003; 144:491–9. doi: 10.1210/en.2002-220734. PMID: 12538609.

9. Kumar TR, Wang Y, Lu N, Matzuk MM. Follicle stimulating hormone is required for ovarian follicle maturation but not male fertility. Nat Genet. 1997; 15:201–4. doi: 10.1038/ng0297-201. PMID: 9020850.

10. Simoni M, Gromoll J, Nieschlag E. The follicle-stimulating hormone receptor: biochemistry, molecular biology, physiology, and pathophysiology. Endocr Rev. 1997; 18:739–73. doi: 10.1210/edrv.18.6.0320. PMID: 9408742.

11. Erickson GF, Hsueh A. Stimulation of aromatase activity by follicle stimulating hormone in rat granulosa cells in vivo and in vitro. Endocrinology. 1978; 102:1275–82. doi: 10.1210/endo-102-4-1275. PMID: 744025.

12. Hillier SG. Gonadotropic control of ovarian follicular growth and development. Mol Cell Endocrinol. 2001; 179:39–46. doi: 10.1016/S0303-7207(01)00469-5. PMID: 11420129.

13. Choi J, Smitz J. Luteinizing hormone and human chorionic gonadotropin: Origins of difference. Mol Cell Endocrinol. 2014; 383:203–13. doi: 10.1016/j.mce.2013.12.009. PMID: 24365330.

14. Ma X, Dong Y, Matzuk MM, Kumar TR. Targeted disruption of luteinizing hormone β-subunit leads to hypogonadism, defects in gonadal steroidogenesis, and infertility. Proc Natl Acad Sci U S A. 2004; 101:17294–9. doi: 10.1073/pnas.0404743101. PMID: 15569941.

15. Themmen AP, Huhtaniemi IT. Mutations of gonadotropins and gonadotropin receptors: elucidating the physiology and pathophysiology of pituitary-gonadal function. Endocr Rev. 2000; 21:551–83. doi: 10.1210/edrv.21.5.0409. PMID: 11041448.

16. Ramaswamy S, Weinbauer GF. Endocrine control of spermatogenesis: Role of FSH and LH/ testosterone. Spermatogenesis. 2014; 4:e996025. doi: 10.1080/21565562.2014.996025. PMID: 26413400.

17. Orth JM. The role of follicle-stimulating hormone in controlling Sertoli cell proliferation in testes of fetal rats. Endocrinology. 1984; 115:1248–55. doi: 10.1210/endo-115-4-1248. PMID: 6090096.

18. Abel MH, Wootton AN, Wilkins V, Huhtaniemi I, Knight PG, Charlton HM. The effect of a null mutation in the follicle-stimulating hormone receptor gene on mouse reproduction. Endocrinology. 2000; 141:1795–803. doi: 10.1210/endo.141.5.7456. PMID: 10803590.

19. Krishnamurthy H, Suresh Babu P, Morales CR, Sairam MR. Delay in sexual maturity of the follicle-stimulating hormone receptor knockout male mouse. Biol Reprod. 2001; 65:522–31. doi: 10.1095/biolreprod65.2.522. PMID: 11466221.

20. Burns KH, Yan C, Kumar TR, Matzuk MM. Analysis of ovarian gene expression in follicle-stimulating hormone β knockout mice. Endocrinology. 2001; 142:2742–51. doi: 10.1210/endo.142.7.8279. PMID: 11415992.

21. Dierich A, Sairam MR, Monaco L, Fimia GM, Gansmuller A, LeMeur M, et al. Impairing follicle-stimulating hormone (FSH) signaling in vivo: Targeted disruption of the FSH receptor leads to aberrant gametogenesis and hormonal imbalance. Proc Natl Acad Sci U S A. 1998; 95:13612–7. doi: 10.1073/pnas.95.23.13612. PMID: 9811848.

22. Lei Z, Mishra S, Zou W, Xu B, Foltz M, Li X, et al. Targeted disruption of luteinizing hormone/human chorionic gonadotropin receptor gene. Mol Endocrinol. 2001; 15:184–200. doi: 10.1210/mend.15.1.0586. PMID: 11145749.

23. Zhang F-P, Poutanen M, Wilbertz J, Huhtaniemi I. Normal prenatal but arrested postnatal sexual development of luteinizing hormone receptor knockout (LuRKO) mice. Mol Endocrinol. 2001; 15:172–83. doi: 10.1210/mend.15.1.0582. PMID: 11145748.

24. Burow S, Mizrahi N, Maugars G, von Krogh K, Nourizadeh-Lillabadi R, Hollander-Cohen L, et al. Characterization of gonadotropin receptors Fshr and Lhr in Japanese medaka, *Oryzias latipes*. Gen Comp Endocrinol. 2020; 285:113276. doi: 10.1016/j.ygcen.2019.113276. PMID: 31536722.

25. Oba Y, Hirai T, Yoshiura Y, Yoshikuni M, Kawauchi H, Nagahama Y. The duality of fish gonadotropin receptors: cloning and functional characterization of a second gonadotropin receptor cDNA expressed in the ovary and testis of amago salmon (*Oncorhynchus rhodurus*). Biochem Biophys Res Commun. 1999; 265:366–71. doi: 10.1006/bbrc.1999.1700. PMID: 10558873.

26. Oba Y, Hirai T, Yoshiura Y, Yoshikuni M, Kawauchi H, Nagahama Y. Cloning, functional characterization, and expression of a gonadotropin receptor cDNA in the ovary and testis of amago salmon (*Oncorhynchus rhodurus*). Biochem Biophys Res Commun. 1999; 263:584–90. doi: 10.1006/bbrc.1999.1346. PMID: 10491336.

27. Miwa S, Yan L, Swanson P. Localization of two gonadotropin receptors in the salmon gonad by in vitro ligand autoradiography. Biol Reprod. 1994; 50:629–42. doi: 10.1095/biolreprod50.3.629. PMID: 8167235.

28. Andersson E, Nijenhuis W, Male R, Swanson P, Bogerd J, Taranger GL, et al. Pharmacological characterization, localization and quantification of expression of gonadotropin receptors in Atlantic salmon (*Salmo salar* L.) ovaries. Gen Comp Endocrinol. 2009; 163:329–39. doi: 10.1016/j.ygcen.2009.05.001. PMID: 19442667.

29. Vischer HF, Granneman JC, Linskens MH, Schulz RW, Bogerd J. Both recombinant African catfish LH and FSH are able to activate the African catfish FSH receptor. J Mol Endocrinol. 2003; 31:133–40. doi: 10.1677/jme.0.0310133. PMID: 12914531.

30. Chauvigné F, Verdura S, Mazón MJ, Duncan N, Zanuy S, Gómez A, et al. Follicle-stimulating hormone and luteinizing hormone mediate the androgenic pathway in Leydig cells of an evolutionary advanced teleost. Biol Reprod. 2012; 87:35. doi: 10.1095/biolreprod.112.100784. PMID: 22649073.

31. Hollander-Cohen L, Bohm B, Hausken K, Levavi-Sivan B. Ontogeny of the specificity of gonadotropin receptors and gene expression in carp. Endocr Connect. 2019; 8:1433–46. doi: 10.1530/EC-19-0389. PMID: 31581128.

32. Bogerd J. Selective ligand-binding determinants in catfish and human gonadotropin receptors. Fish Physiol Biochem. 2005; 31:247–54. doi: 10.1007/s10695-006-0032-3. PMID: 20035466.

33. Kwok HF, So WK, Wang Y, Ge W. Zebrafish gonadotropins and their receptors: I. Cloning and characterization of zebrafish follicle-stimulating hormone and luteinizing hormone receptors—evidence for their distinct functions in follicle development. Biol Reprod. 2005; 72:1370–81. doi: 10.1095/biolreprod.104.038190. PMID: 15728795.

34. So WK, Kwok HF, Ge W. Zebrafish gonadotropins and their receptors: II. Cloning and characterization of zebrafish follicle-stimulating hormone and luteinizing hormone subunits—their spatial-temporal expression patterns and receptor specificity. Biol Reprod. 2005; 72:1382–96. doi: 10.1095/biolreprod.104.038216. PMID: 15728794.

35. Zhang Z, Zhu B, Ge W. Genetic analysis of zebrafish gonadotropin (FSH and LH) functions by TALEN-mediated gene disruption. Mol Endocrinol. 2015; 29:76–98. doi: 10.1210/me.2014-1256. PMID: 25396299.

36. Zhang Z, Lau SW, Zhang L, Ge W. Disruption of zebrafish follicle-stimulating hormone receptor (fshr) but not luteinizing hormone receptor (lhcgr) gene by TALEN leads to failed follicle activation in females followed by sexual reversal to males. Endocrinology. 2015; 156:3747–62. doi: 10.1210/en.2015-1039. PMID: 25993524.

37. Chu L, Li J, Liu Y, Cheng CH. Gonadotropin signaling in zebrafish ovary and testis development: insights from gene knockout study. Mol Endocrinol. 2015; 29:1743–58. doi: 10.1210/me.2015-1126. PMID: 26452104.

38. Rosenbaum DM, Rasmussen SG, Kobilka BK. The structure and function of G-protein-coupled receptors. Nature. 2009; 459:356–63. doi: 10.1038/nature08144. PMID: 19458711

39. Milligan G. Constitutive activity and inverse agonists of G protein-coupled receptors: a current perspective. Mol Pharmacol. 2003; 64:1271–6. doi: 10.1124/mol.64.6.1271. PMID: 14645655.

40. Baker PJ, Pakarinen P, Huhtaniemi IT, Abel MH, Charlton HM, Kumar TR, et al. Failure of normal Leydig cell development in follicle-stimulating hormone (FSH) receptor-deficient mice, but not FSHβ-deficient mice: role for constitutive FSH receptor activity. Endocrinology. 2003; 144:138–45. doi: 10.1210/en.2002-220637. PMID: 12488339.

41. Kumar RS, Ijiri S, Trant JM. Molecular biology of the channel catfish gonadotropin receptors: 2. Complementary DNA cloning, functional expression, and seasonal gene expression of the follicle-stimulating hormone receptor. Biol Reprod. 2001; 65:710–7. doi: 10.1095/biolreprod65.3.710. PMID: 11514332.

42. Vischer H, Bogerd J. Cloning and functional characterization of a gonadal luteinizing hormone receptor complementary DNA from the African catfish (*Clarias gariepinus*). Biol Reprod. 2003; 68:262–71. doi: 10.1095/biolreprod.102.004515. PMID: 12493722.

43. Meeker ND, Hutchinson SA, Ho L, Trede NS. Method for isolation of PCR-Ready genomic DNA from zebrafish tissues. Biotechniques. 2007; 43:610–4. doi: 10.2144/000112619. PMID: 18072590.

44. Zhai Y, Zhang X, Zhao C, Geng R, Wu K, Yuan M, et al. Rescue of bmp15 deficiency in zebrafish by mutation of inha reveals mechanisms of BMP15 regulation of folliculogenesis. PLoS Genet. 2023; 19:e1010954. doi: 10.1371/journal.pgen.1010954. PMID: 37713421.

45. Zhou R, Tsang AH, Lau SW, Ge W. Pituitary adenylate cyclase-activating polypeptide (PACAP) and its receptors in the zebrafish ovary: evidence for potentially dual roles of PACAP in controlling final oocyte maturation. Biol Reprod. 2011; 85:615–25. doi: 10.1095/biolreprod.111.091884. PMID: 21636738.

46. Song W, Lu H, Wu K, Zhang Z, Lau ESW, Ge W. Genetic evidence for estrogenicity of bisphenol A in zebrafish gonadal differentiation and its signalling mechanism. J Hazard Mater. 2020; 386:121886. doi: 10.1016/j.jhazmat.2019.121886. PMID: 31887561.

47. Shang G, Peng X, Ji C, Zhai G, Ruan Y, Lou Q, et al. Steroidogenic acute regulatory protein and luteinizing hormone are required for normal ovarian steroidogenesis and oocyte maturation in zebrafish. Biol Reprod. 2019; 101:760–70. doi: 10.1093/biolre/ioz132. PMID: 31322169.

48. Chu L, Li J, Liu Y, Hu W, Cheng CH. Targeted gene disruption in zebrafish reveals noncanonical functions of LH signaling in reproduction. Mol Endocrinol. 2014; 28:1785–95. doi: 10.1210/me.2014-1061. PMID: 25238195.

49. Liu Y, Kossack ME, McFaul ME, Christensen LN, Siebert S, Wyatt SR, et al. Single-cell transcriptome reveals insights into the development and function of the zebrafish ovary. eLife. 2022; 11:e76014. doi: 10.7554/eLife.76014. PMID: 35588359.

50. Bogerd J. Ligand-selective determinants in gonadotropin receptors. Mol Cell Endocrinol. 2007; 260-262:144–52. doi: 10.1016/j.mce.2006.01.019. PMID: 17055148.

51. Xie Y, Chu L, Liu Y, Sham KWY, Li J, Cheng CHK. The highly overlapping actions of Lh signaling and Fsh signaling on zebrafish spermatogenesis. J Endocrinol. 2017; 234:233–46. doi: 10.1530/joe-17-0079. PMID: 28611209.

52. Li J, Cheng CHK. Evolution of gonadotropin signaling on gonad development: insights from gene knockout studies in zebrafish. Biol Reprod. 2018; 99:686–94. doi: 10.1093/biolre/ioy101. PMID: 29718109.

53. Kleinau G, Biebermann H. Constitutive activities in the thyrotropin receptor: regulation and significance. Adv Pharmacol. 2014; 70:81–119. doi: 10.1016/B978-0-12-417197-8.00003-1. PMID: 24931193.

54. Seifert R, Wenzel-Seifert K. Constitutive activity of G-protein-coupled receptors: cause of disease and common property of wild-type receptors. Naunyn Schmiedebergs Arch Pharmacol. 2002; 366:381–416. doi: 10.1007/s00210-002-0588-0. PMID: 12382069.

55. Sola L, Gornung E. Classical and molecular cytogenetics of the zebrafish, *Danio rerio* (Cyprinidae, Cypriniformes): an overview. Genetica. 2001; 111:397–412. doi: 10.1023/A:1013776323077. PMID: 11841183.

56. Kashimada K, Koopman P. Sry: the master switch in mammalian sex determination. Development. 2010; 137:3921–30. doi: 10.1242/dev.048983. PMID: 21062860.

57. Matsuda M, Nagahama Y, Shinomiya A, Sato T, Matsuda C, Kobayashi T, et al. DMY is a Y-specific DM-domain gene required for male development in the medaka fish. Nature. 2002; 417:559–63. doi: 10.1038/nature751. PMID: 12037570.

58. Liew WC, Bartfai R, Lim Z, Sreenivasan R, Siegfried KR, Orban L. Polygenic sex determination system in zebrafish. PLoS One. 2012; 7:e34397. doi: 10.1371/journal.pone.0034397. PMID: 22506019.

59. Shang EH, Yu RM, Wu RS. Hypoxia affects sex differentiation and development, leading to a male-dominated population in zebrafish (*Danio rerio*). Environ Sci Technol. 2006; 40:3118–22. doi: 10.1021/es0522579. PMID: 16719120.

60. Valdivieso A, Caballero-Huertas M, Moraleda-Prados J, Piferrer F, Ribas L. Exploring the effects of rearing densities on epigenetic modifications in the zebrafish gonads. Int J Mol Sci. 2023; 24:16002. doi: 10.3390/ijms242116002. PMID: 37958987.

61. Uchida D, Yamashita M, Kitano T, Iguchi T. An aromatase inhibitor or high water temperature induce oocyte apoptosis and depletion of P450 aromatase activity in the gonads of genetic female zebrafish during sex-reversal. Comp Biochem Physiol A Physiol. 2004; 137:11–20. doi: 10.1016/S1095-6433(03)00178-8. PMID: 14720586.

62. Lawrence C, Ebersole JP, Kesseli RV. Rapid growth and out-crossing promote female development in zebrafish (*Danio rerio*). Environ Biol Fishes. 2008; 81:239–46. doi: 10.1007/s10641-007-9195-8.

63. Dranow DB, Tucker RP, Draper BW. Germ cells are required to maintain a stable sexual phenotype in adult zebrafish. Dev Biol. 2013; 376:43–50. doi: 10.1016/j.ydbio.2013.01.016. PMID: 23348677.

64. Kobayashi Y, Nakamura M, Sunobe T, Usami T, Kobayashi T, Manabe H, et al. Sex change in the Gobiid fish is mediated through rapid switching of gonadotropin receptors from ovarian to testicular portion or vice versa. Endocrinology. 2009; 150:1503–11. doi: 10.1210/en.2008-0569. PMID: 18948407.

65. Yamaguchi T, Yamaguchi S, Hirai T, Kitano T. Follicle-stimulating hormone signaling and Foxl2 are involved in transcriptional regulation of aromatase gene during gonadal sex differentiation in Japanese flounder, *Paralichthys olivaceus*. Biochem Biophys Res Commun. 2007; 359:935–40. doi: 10.1016/j.bbrc.2007.05.208. PMID: 17574208.

66. Takahashi A, Kanda S, Abe T, Oka Y. Evolution of the hypothalamic-pituitary-gonadal axis regulation in vertebrates revealed by knockout medaka. Endocrinology. 2016; 157:3994–4002. doi: 10.1210/en.2016-1356. PMID: 27560548.

67. Moses E, Franek R, Harel I. A scalable and tunable platform for functional interrogation of peptide hormones in fish. eLife. 2023; 12. doi: 10.7554/eLife.85960. PMID: 37872843.

68. Murozumi N, Nakashima R, Hirai T, Kamei Y, Ishikawa-Fujiwara T, Todo T, et al. Loss of follicle-stimulating hormone receptor function causes masculinization and suppression of ovarian development in genetically female medaka. Endocrinology. 2014; 155:3136–45. doi: 10.1210/en.2013-2060. PMID: 24877625.

69. Andersson E, Schulz RW, Almeida F, Kleppe L, Skaftnesmo KO, Kjaerner-Semb E, et al. Loss of Fshr prevents testicular maturation in Atlantic salmon (*Salmo salar* L.). Endocrinology. 2024; 165:bqae013. doi: 10.1210/endocr/bqae013. PMID: 38298132.

70. Leal M, Skaar K, França L, Schulz R. The effects of IGF-I on zebrafish testis in tissue culture. Anim Reprod. 2006; 3:181.

71. Zhang Z, Wu K, Ren Z, Ge W. Genetic evidence for Amh modulation of gonadotropin actions to control gonadal homeostasis and gametogenesis in zebrafish and its noncanonical signaling through Bmpr2a receptor. Development. 2020; 147:dev189811. doi: 10.1242/dev.189811. PMID: 33060133.

72. Zhang Z, Zhu B, Chen W, Ge W. Anti-Müllerian hormone (Amh/amh) plays dual roles in maintaining gonadal homeostasis and gametogenesis in zebrafish. Mol Cell Endocrinol. 2020; 517:110963. doi: 10.1016/j.mce.2020.110963. PMID: 32745576.

73. Nagi K, Onaran HO. Biased agonism at G protein-coupled receptors. Cell Signal. 2021; 83:109981. doi: 10.1016/j.cellsig.2021.109981. PMID: 33744417.

74. Wootten D, Christopoulos A, Marti-Solano M, Babu MM, Sexton PM. Mechanisms of signalling and biased agonism in G protein-coupled receptors. Nat Rev Mol Cell Biol. 2018; 19:638–53. doi: 10.1038/s41580-018-0049-3. PMID: 30104700.

75. Riccetti L, Yvinec R, Klett D, Gallay N, Combarnous Y, Reiter E, et al. Human luteinizing hormone and chorionic gonadotropin display biased agonism at the LH and LH/CG receptors. Sci Rep. 2017; 7:940. doi: 10.1038/s41598-017-01078-8. PMID: 28424471.

76. Zariñán T, Butnev VY, Gutiérrez-Sagal R, Maravillas-Montero JL, Martínez-Luis I, Mejía-Domínguez NR, et al. In vitro impact of FSH glycosylation variants on FSH receptor-stimulated signal transduction and functional selectivity. J Endocr Soc. 2020; 4:bvaa019. doi: 10.1210/jendso/bvaa019. PMID: 32342021.

77. Miller WL, Auchus RJ. The molecular biology, biochemistry, and physiology of human steroidogenesis and its disorders. Endocr Rev. 2011; 32:81–151. doi: 10.1210/er.2010-0013. PMID: 21051590.

78. Drummond AE. The role of steroids in follicular growth. Reprod Biol Endocrinol. 2006; 4:1–11. doi: 10.1186/1477-7827-4-16. PMID: 16603089.

79. Ogiwara K, Takahashi T. Involvement of the nuclear progestin receptor in LH-induced expression of membrane type 2-matrix metalloproteinase required for follicle rupture during ovulation in the medaka, *Oryzias latipes*. Mol Cell Endocrinol. 2017; 450:54–63. doi: 10.1016/j.mce.2017.04.016. PMID: 28416325.

80. Tokumoto T, Yamaguchi T, Ii S, Tokumoto M. In vivo induction of oocyte maturation and ovulation in zebrafish. PLoS One. 2011; 6:e25206. doi: 10.1371/journal.pone.0025206. PMID: 21980399.

81. Shi S, Zhang Y, Huang J, Lou Q, Jin X, He J, et al. Effective “off-on” switch for fertility control in female zebrafish. Front Mar Sci. 2024; 11:1381305. doi: 10.3389/fmars.2024.1381305.

82. Hagiwara A, Ogiwara K, Katsu Y, Takahashi T. Luteinizing hormone-induced expression of Ptger4b, a prostaglandin E2 receptor indispensable for ovulation of the Medaka *Oryzias latipes*, is regulated by a genomic mechanism involving nuclear progestin receptor. Biol Reprod. 2014; 90:126. doi: 10.1095/biolreprod.113.115485. PMID: 24790162.

83. Zhu Y, Liu D, Shaner ZC, Chen S, Hong W, Stellwag EJ. Nuclear progestin receptor (pgr) knockouts in zebrafish demonstrate role for pgr in ovulation but not in rapid non-genomic steroid mediated meiosis resumption. Front Endocrinol. 2015; 6:37. doi: 10.3389/fendo.2015.00037. PMID: 25852646.

84. Wu X, Liu D, Chen S, Hong W, Zhu Y. Impaired oocyte maturation and ovulation in membrane progestin receptor (mPR) knockouts in zebrafish. Mol Cell Endocrinol. 2020; 511:110856. doi: 10.1016/j.mce.2020.110856. PMID: 32387526.

85. Liu DT, Carter NJ, Wu XJ, Hong WS, Chen SX, Zhu Y. Progestin and nuclear progestin receptor are essential for upregulation of metalloproteinase in zebrafish preovulatory follicles. Front Endocrinol. 2018; 9:517. doi: 10.3389/fendo.2018.00517. PMID: 30279677.

86. Liu DT, Hong WS, Chen SX, Zhu Y. Upregulation of adamts9 by gonadotropin in preovulatory follicles of zebrafish. Mol Cell Endocrinol. 2020; 499:110608. doi: 10.1016/j.mce.2019.110608. PMID: 31586455.

87. Tang H, Liu Y, Li J, Yin Y, Li G, Chen Y, et al. Gene knockout of nuclear progesterone receptor provides insights into the regulation of ovulation by LH signaling in zebrafish. Sci Rep. 2016; 6:28545. doi: 10.1038/srep28545. PMID: 27333837.

88. Park JY, Su YQ, Ariga M, Law E, Jin S-LC, Conti M. EGF-like growth factors as mediators of LH action in the ovulatory follicle. Science. 2004; 303:682–4. doi: 10.1126/science.1092463. PMID: 14726596.

89. Richani D, Gilchrist RB. The epidermal growth factor network: role in oocyte growth, maturation and developmental competence. Hum Reprod Update. 2018; 24:1–14. doi: 10.1093/humupd/dmx029. PMID: 29029246.

90. Wang Y, Ge W. Cloning of epidermal growth factor (EGF) and EGF receptor from the zebrafish ovary: evidence for EGF as a potential paracrine factor from the oocyte to regulate activin/follistatin system in the follicle cells. Biol Reprod. 2004; 71:749–60. doi: 10.1095/biolreprod.104.028399. PMID: 15115721.

91. Tse AC, Ge W. Spatial localization of EGF family ligands and receptors in the zebrafish ovarian follicle and their expression profiles during folliculogenesis. Gen Comp Endocrinol. 2010; 167:397–407. doi: 10.1016/j.ygcen.2009.09.012. PMID: 19799903.

92. Song Y, Chen W, Zhu B, Ge W. Disruption of epidermal growth factor receptor but not EGF blocks follicle activation in zebrafish ovary. Front Cell Dev Biol. 2021; 9:750888. doi: 10.3389/fcell.2021.750888. PMID: 35111746.

93. Pang Y, Ge W. Epidermal growth factor and TGFα promote zebrafish oocyte maturation *in vitro*: potential role of the ovarian activin regulatory system. Endocrinology. 2002; 143:47–54. doi: 10.1210/endo.143.1.8579. PMID: 11751590.

94. Chung CK, Ge W. Epidermal growth factor differentially regulates activin subunits in the zebrafish ovarian follicle cells via diverse signaling pathways. Mol Cell Endocrinol. 2012; 361:133–42. doi: 10.1016/j.mce.2012.03.022. PMID: 22503865.

95. Chung CK, Ge W. Human chorionic gonadotropin (hCG) induces MAPK3/1 phosphorylation in the zebrafish ovarian follicle cells independent of EGF/EGFR pathway. Gen Comp Endocrinol. 2013; 188:251–7. doi: 10.1016/j.ygcen.2013.04.020. PMID: 23644153.

96. Pang Y, Ge W. Activin stimulation of zebrafish oocyte maturation in vitro and its potential role in mediating gonadotropin-induced oocyte maturation. Biol Reprod. 1999; 61:987–92. doi: 10.1095/biolreprod61.4.987. PMID: 10491634.

97. Pang Y, Ge W. Gonadotropin and activin enhance maturational competence of oocytes in the zebrafish (*Danio rerio*). Biol Reprod. 2002; 66:259–65. doi: 10.1095/biolreprod66.2.259. PMID: 11804937.

98. Petrino TR, Toussaint G, Lin YWP. Role of inhibin and activin in the modulation of gonadotropin-and steroid-induced oocyte maturation in the teleost Fundulus heteroclitus. Reprod Biol Endocrinol. 2007; 5:1–9. doi: 10.1186/1477-7827-5-21. PMID: 17550604.

99. Ge W. Activin and its receptors in fish reproduction. Hormones and Their Receptors in Fish Reproduction. 2005:128–54. doi: 10.1142/9789812569189_0005.

100. Wu T, Patel H, Mukai S, Melino C, Garg R, Ni X, et al. Activin, inhibin, and follistatin in zebrafish ovary: expression and role in oocyte maturation. Biol Reprod. 2000; 62:1585–92. doi: 10.1095/biolreprod62.6.1585. PMID: 10819759.

